# Prepronociceptin expressing neurons in the extended amygdala encode and promote rapid arousal responses to motivationally salient stimuli

**DOI:** 10.1101/2020.01.21.914341

**Authors:** Jose Rodriguez-Romaguera, Randall L Ung, Hiroshi Nomura, James M Otis, Marcus L Basiri, Vijay MK Namboodiri, Xueqi Zhu, J Elliott Robinson, Jenna A McHenry, Oksana Kosyk, Thomas C Jhou, Thomas L Kash, Michael R Bruchas, Garret D Stuber

## Abstract

Motivational states are complex and consist of cognitive, emotional, and physiological components controlled by a network across multiple brain regions. An integral component of this neural circuitry is the bed nucleus of the stria terminalis (BNST). Here, we identified a subpopulation of neurons within BNST expressing the gene prepronociceptin (*Pnoc*^BNST^), that can modulate the rapid changes in physiological arousal that occur upon exposure to stimuli with motivational salience. Using *in vivo* two-photon calcium imaging we found that excitatory responses from individual *Pnoc*^BNST^ neurons directly corresponded with rapid increases in pupillary size and occurred upon exposure to both aversive and rewarding odors. Furthermore, optogenetic activation of these neurons increased pupillary size, but did not alter approach/avoidance or locomotor behaviors. These findings suggest that excitatory responses in *Pnoc*^BNST^ neurons encode rapid arousal responses irrespective of tested behaviors. Further histological, electrophysiological, and single-cell RNA sequencing data revealed that *Pnoc*^BNST^ neurons are composed of genetically and anatomically identifiable subpopulations that can be further investigated. Taken together, our findings demonstrate a key role for a *Pnoc*^BNST^ neuronal ensemble in encoding the rapid arousal responses that are triggered by motivational stimuli.

## INTRODUCTION

Dysfunctional arousal responses are a core component of many neuropsychiatric disorders. For example, patients with anxiety disorders often show hyperarousal responses to negatively salient stimuli, and patients suffering from depression show hypoarousal responses to positively salient stimuli (Craske et al., 2009; Lang and McTeague, 2009; Patriquin et al., 2019; Schmidt et al., 2017; Urbano et al., 2017; Wilhelm and Roth, 2001). Elucidating the neural circuit elements that orchestrate changes in physiological arousal are thus essential for understanding maladaptive motivational states (Marton and Sohal, 2016; Sparta et al., 2013; Touriño et al., 2013). Rodent models allow for the dissection of the neural circuits for both negative and positive motivational states by presenting stimuli that elicit aversion or reward (Calhoon and Tye, 2015; Stuber and Wise, 2016; Tovote et al., 2015). However, these studies often overlook the rapid increases in physiological arousal that characterize changing motivational states. In humans, rapid (within seconds) increases in physiological arousal, as measured by pupil size changes, follow exposure to negatively salient stimuli, such as threat-inducing images (Cascardi et al., 2015; Price et al., 2013). The same is true when humans are presented with positively salient stimuli, such as rewarding images of money or videos of caregivers (Schneider et al., 2018; Tummeltshammer et al., 2019). Thus, in addition to long-term adaptations in arousal (e.g. sleep/wake states (de Lecea et al., 2012)), an important component of motivation are these rapid changes in physiological arousal upon presentation of salient stimuli.

Evidence from anatomical (Dabrowska et al., 2011; Dong et al., 2001; Singewald et al., 2003), behavioral (Duvarci et al., 2009; Jennings et al., 2013a; Kim et al., 2013; Walker et al., 2009), and neuroimaging (Straube et al., 2007; Yassa et al., 2012) studies have implicated the bed nucleus of the stria terminalis (BNST, a part of the extended amygdala) as a key component of the neural circuitry that regulates motivated behavior. Historically, the role of the BNST in physiological arousal has been largely overlooked, although this region is involved in a variety of motivational states. For instance, recent neurocircuit studies in mice have highlighted the role of BNST in reward and aversion (Giardino et al., 2018; Jennings et al., 2013a, 2013b; Kim et al., 2013), fear and anxiety-like behaviors (Crowley et al., 2016; Duvarci et al., 2009; Kim et al., 2013; Marcinkiewcz et al., 2016; Walker et al., 2009), and social preference and aversion (Goodson and Wang, 2006; Lei et al., 2010; Newman Sarah Winans, 2006). Further, previous studies have identified how subsets of BNST neurons expressing certain marker genes such as corticotrophin-releasing hormone (*Crh*^BNST^), protein kinase C δ (*Pkcδ* ^BNST^), and somatostatin (*Som*^BNST^) drive motivated behaviors (Kash et al., 2015; Koob and Heinrichs, 1999; Lebow and Chen, 2016; Tovote et al., 2015). However, whether specific neural populations within the BNST drive rapid changes in physiological arousal remains unknown. This is in part due to the low number of studies linking the functional heterogeneity within BNST with its role in rapid changes in physiological arousal (Kim et al., 2013).

Neuropeptide gene expression patterns have identified functionally distinct subpopulations of neurons in BNST (Kash et al., 2015). Recently, neurons that express the prepronociceptin gene (*Pnoc*, the genetic precursor to the nociception neuropeptide) within the central nucleus of the amygdala and the paranigral ventral tegmental area were shown to have a role in gating motivational states and reward seeking (Hardaway et al., 2019; Parker et al., 2019). Since the BNST contains many neurons that express the *Pnoc* gene (Boom et al., 1999; Ikeda et al., 1998), we set out to investigate the role of these neurons in contributing towards aspects of motivational states, in particular in driving the physiological arousal responses that occurs in response to motivationally salient stimuli.

In the present study, we used cell-type-specific optogenetic and head-fixed two-photon calcium imaging approaches (McHenry et al., 2017; Namboodiri et al., 2019; Otis et al., 2017) to assess the role of *Pnoc*^BNST^ neurons in driving and encoding rapid physiological arousal responses to aversive and rewarding odors. We then characterized the anatomical connectivity and genetic identity of *Pnoc*^BNST^ neurons using a combination of histological, electrophysiological, and single-cell sequencing approaches. *In vivo* calcium imaging revealed heterogeneous response dynamics in *Pnoc*^BNST^ neurons. However, individual cells that encoded rapid changes in physiological arousal (as measured by pupillary dynamics) showed excitatory responses when mice where exposed to either aversive or rewarding odors. We also found that optogenetic activation of *Pnoc*^BNST^ neurons did not induce approach/avoidance or locomotor behaviors, but specifically increased physiological measurements associated with arousal (pupillary response and heart rate). scRNAseq revealed that *Pnoc*^BNST^ neurons are transcriptionally diverse and can be further subdivided by multiple distinct gene markers. Collectively, these results suggest that *Pnoc*^BNST^ neurons are molecularly heterogeneous, but that they play an important role in orchestrating arousal-related responses associated with motivationally salient stimuli.

## RESULTS

### Expression of prepronociceptin defines a subpopulation of GABAergic neurons within the adBNST that can be monitored using calcium imaging approaches

The BNST is composed of various subnuclei that have unique molecular and functional identities (Giardino et al., 2018; Gungor and Paré, 2016). Therefore, we first assessed the distribution of *Pnoc*-expressing neurons across the BNST. Using fluorescent *in situ* hybridization (FISH), we observed that *Pnoc*^BNST^ neurons were distributed throughout the BNST (Figure 1A) but enriched in the anterodorsal BNST (adBNST), as previously described (Neal et al., 1999). Further, we found that *Pnoc*^BNST^ neurons predominantly express the vesicular GABA transporter gene, *Slc32a1*, (*Vgat)* and not the vesicular glutamate transporter 2 gene, *Slc17a6*, (*Vglut2*; Figure 1B), indicating that *Pnoc* expression defines a subpopulation of GABAergic neurons within BNST. *Pnoc*-IRES-*Cre* mice (Hardaway et al., 2019; Parker et al., 2019) were then used for selective targeting of *Pnoc^+^* neurons in the adBNST in conjunction with cre-inducible viruses.

**Figure 1.**
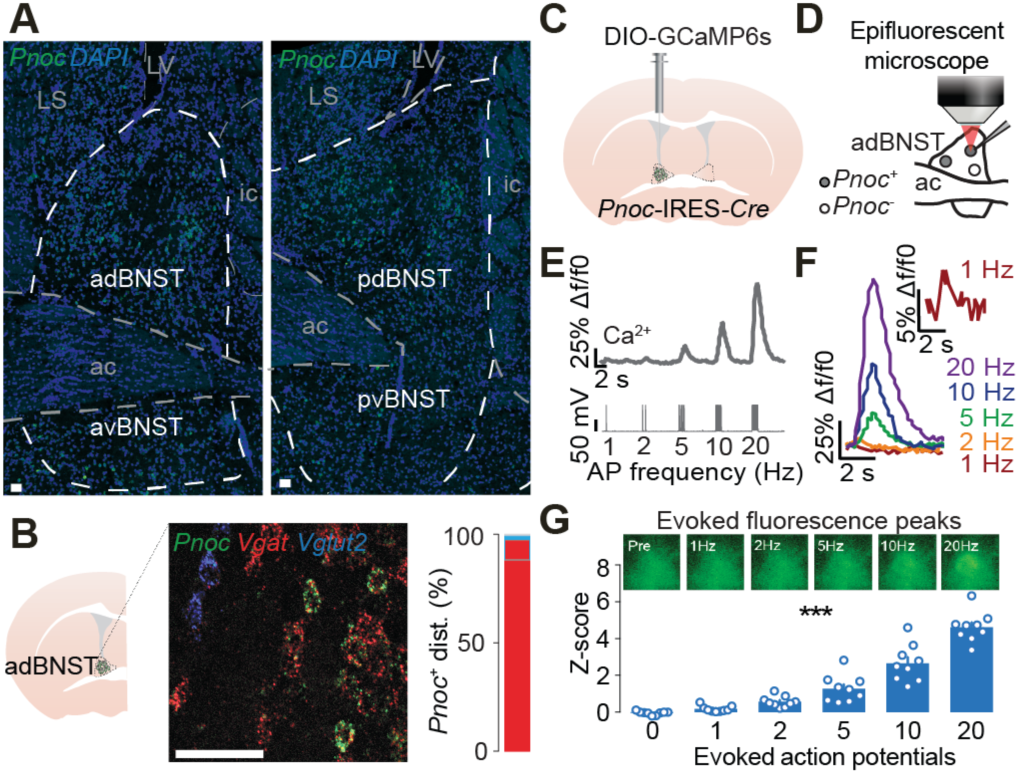
(**A**) Confocal images depicting the distribution of *Pnoc*-expressing neurons in BNST using FISH with DAPI counterstain. Abbreviations: LV = lateral ventricle; LS = lateral septum; adBNST = anterodorsal BNST; avBNST = anteroventral BNST; pdBNST = posterodorsal BNST; pvBNST = posteroventral BNST; ac = anterior commissure. (**B**) Confocal image depicting the overlap between the expression of *Pnoc*, *Vgat*, and *Vglut2* within BNST neurons using FISH (left). Proportion of *Vgat^+^* and *Vglut2^+^* neurons quantified using FISH (right). (**C**) Schematic of injection of AAVdj-EF1α-DIO-GCaMP6s into the adBNST of *Pnoc*-IRES-*Cre* mice. (**D**) Schematic of simultaneous patch-clamp electrophysiology and calcium imaging of GCaMP6s-expressing *Pnoc*^+^ neurons. (**E**) Sample traces showing a series of depolarizing pulses (1–20 Hz) applied in current-clamp mode to drive trains of action potentials (bottom), during which GCaMP6s fluorescence was tracked in recorded neurons (top). (**F**) Overlay of sample traces showing elevation of GCaMP6s fluorescence signal during the depolarizing pulses, such that a single action potential was detectable (red waveform). (**G**) Representative images of an individual *Pnoc*^BNST^ neuron showing evoked fluorescence peaks at the various depolarizing pulses (top). Action potential generation resulted in linear elevations in GCaMP6s fluorescence (bottom). Data shown as mean ± SEM. \*\*\**p<0.001*.

To characterize how *Pnoc*^BNST^ firing related to calcium mediated fluorescent dynamics, we transduced the adBNST of *Pnoc*-Cre mice with Cre-dependent GCaMP6s virus (Figure 1C). We then performed calcium imaging under an epifluorescent microscope and simultaneously activated *Pnoc*^BNST^ neurons via current injections at various frequencies. We found a linear relationship between evoked action potentials and their respective fluorescent peaks (Figure 1E-G), demonstrating that calcium dynamics track evoked firing in brain slices.

### *Pnoc*^BNST^ neurons encode rapid changes in arousal to aversive and rewarding stimuli

Since the BNST is thought to coordinate motivational states essential for guiding actions of reward seeking and aversion (Calhoon and Tye, 2015; Kash et al., 2015; Lebow and Chen, 2016; Stamatakis et al., 2014; Tovote et al., 2015), we tested if activity of *Pnoc*^BNST^ neurons is altered by exposure to stimuli with opposing motivational salience. We exposed mice to either trimethylthiazoline (TMT, as an aversive odor) or peanut oil (as an appetitive odor), odors that induce either place aversion or place preference (Root et al., 2014), respectively. First, we demonstrated that freely moving mice reliably avoided a location with TMT and preferred a location containing peanut oil, (Figure 2A-C) confirming the aversive and appetitive nature of these olfactory stimuli. Pupillary responses have been shown to reflect rapid changes in physiological arousal (Cascardi et al., 2015; Price et al., 2013). Consistent with this idea, we also observed increases in pupillary size when freely-moving mice were in close proximity to either TMT or peanut containing odor swabs, as compared to a swab with water as a control (Figure 2D-F). To evaluate encoding of *Pnoc*^BNST^ neurons to these odors, we developed a head-fixed behavioral preparation compatible with two-photon microscopy to control proximity of an odor swab and allow us to measure pupillometry and ambulation (Figure 2J). Odors where presented with a cotton swab that could be positioned near or far from the mouse while pupil size was recorded through a camera aimed at one of the eyes (Reimer et al., 2014). Having animals head-fixed also allowed us to record calcium activity from individual *Pnoc*^BNST^ neurons in live awake mice via a GRIN lens under a two-photon microscope (Figures 2G-H). Using algorithms that employ constrained non-negative matrix factorization (CNMF) we extracted traces of activity dynamics from individual *Pnoc*^BNST^ neurons (Figures 2I; Figure S1A-E).

**Figure 2.**
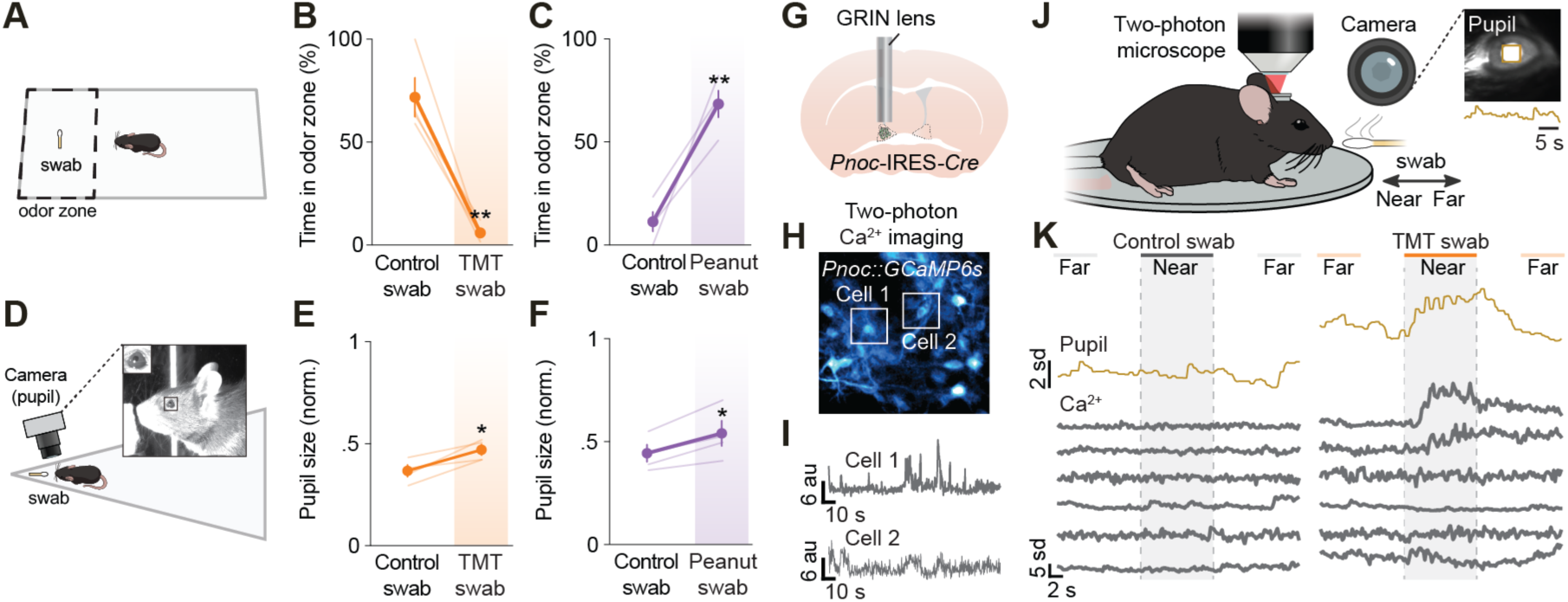
(**A**) Schematic of a freely moving mouse in its home cage to assess avoidance and approach behaviors to a water control, TMT, or peanut oil containing swabs. (**B**) Group average of time in odor zone in mice exposed consecutively to control and TMT swabs (swab was placed in *preferred side* determined during a baseline prior to testing). (**C**) Group average of time in odor zone in mice exposed consecutively to control and peanut swabs (swab was placed in *non-preferred* side determined during a baseline prior to testing). (**D**) Schematic of modified arena used for pupillometry in freely moving animals during exposure to control, TMT, or peanut containing swabs. Image of a mouse sniffing the odor swab. Insert: Close up of the mouse’s pupil. (**E**) Group average of normalized pupil diameter between consecutive exposure to control and TMT odor during the first initial contact with the odor swab. (**F**) Group average of normalized pupil diameter between consecutive exposure to control and peanut odor during the first initial contact with the odor swab. (**G**) Schematic of implantation of a GRIN lens above adBNST of *Pnoc*-IRES-*Cre* mice injected with AAVdj-EF1α-DIO-GCaMP6s. (**H**) Representative image of *Pnoc*^BNST^ neurons through a GRIN lens. (**I**) Extracted calcium traces from two representative *Pnoc*^BNST^ neurons using CNMF. (**J**) Schematic of a head-fixed mouse on a running disc with simultaneous pupillometry under a two-photon microscope while being exposed to a movable odor swab. The odor swab was either 25 cm (far) or 1 cm (near) from the mice. *Inset*: Representative frame of a mouse pupil with size tracking square and accompanying sample pupil trace. (**K**) Sample traces of *Pnoc*^BNST^ neurons shown based on location of either the control or TMT swab. Data shown as mean ± SEM. \**p<0.05,* \*\**p<0.01*.

We found that over 50% of *Pnoc*^BNST^ neurons showed a significant change in response (either excitation or inhibition) to a swab with either odorant, as compared to a control swab (Figure 3A-F). Strikingly, neurons that showed significant excitation or inhibition to TMT or peanut oil swabs showed significant correlations between pupillary fluctuations and their individual neural dynamics, whereas the control swab did not (Figure 3G-K). We also found different proportion of neurons that were excited and inhibited between TMT and peanut odor exposure (Figures 3G, 3J), suggesting that subtypes of *Pnoc*^BNST^ neurons may exist to encode aversive vs. rewarding arousal states. The subpopulation of neurons that showed significant inhibitory responses appears to be specific to aversive arousal states. Excitatory responses were observed in both aversive and rewarding arousal states, with a larger proportion of cells showing significant excitation to rewarding arousal states (Figures 3G, 3J). Since both aversive and rewarding arousal states induced an increase in excitatory neuronal responses that correlated with pupillary dynamics, our data suggest that excitatory responses of *Pnoc*^BNST^ neurons encode increases in arousal.

**Figure 3.**
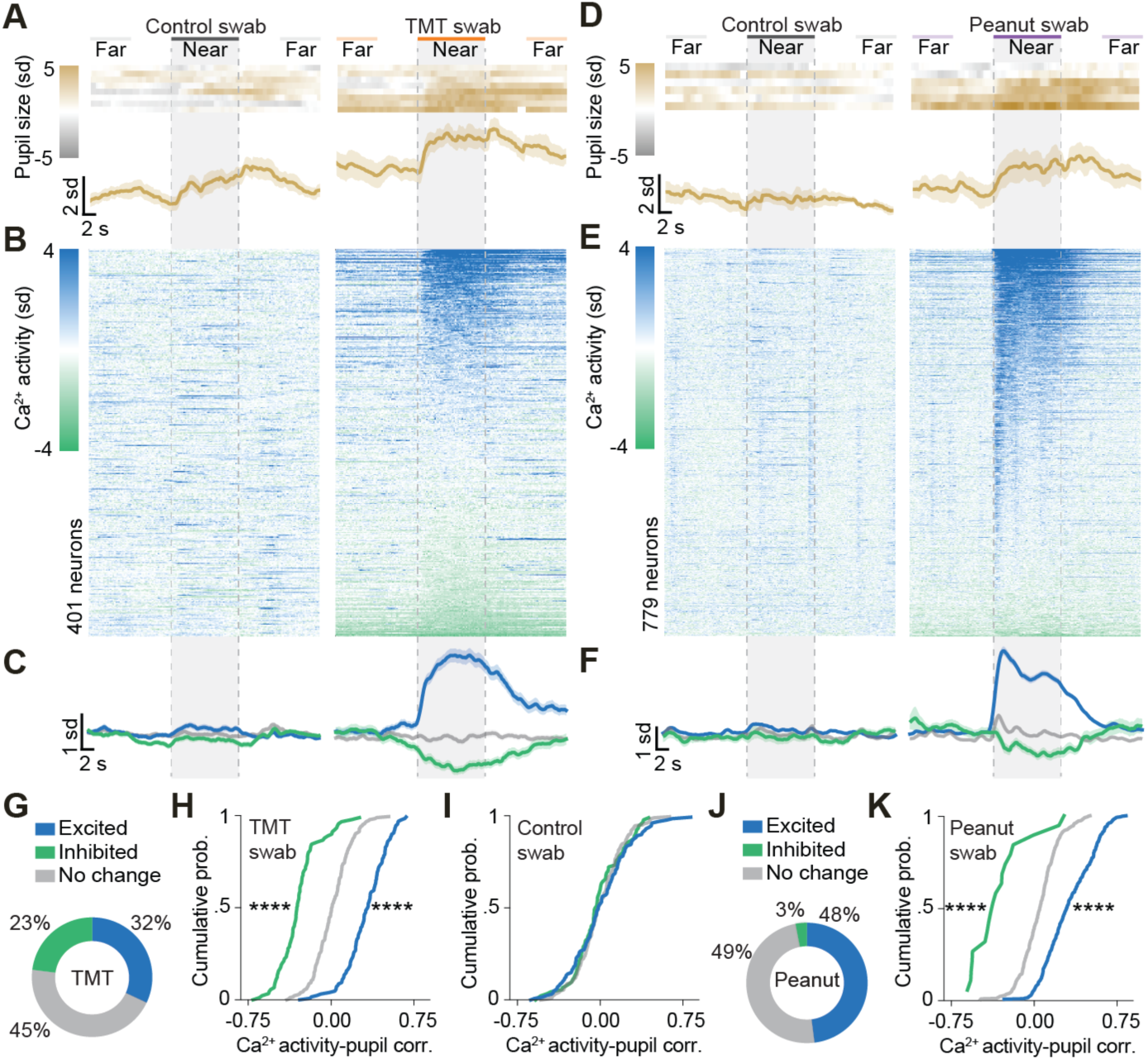
(**A**) Heat map of individual data (top) and average group data (bottom) for pupil responses to control and TMT swab. (**B**) Heat map of responses to the control and TMT swabs from all *Pnoc*^BNST^ neurons, organized by their average response to the TMT swab. (**C**) Response dynamics of *Pnoc*^BNST^ neurons to the control and TMT swabs that showed significant excitatory, inhibitory, or no change in activity to the TMT swab. (**D**) Heat map of individual data (top) and average group data (bottom) for pupil responses to control and peanut swab. (**E**) Heat map of responses to the control and peanut swabs from all *Pnoc*^BNST^ neurons, organized by their average response to the peanut swab. (**F**) Response dynamics of *Pnoc*^BNST^ neurons to the control and peanut swabs that showed significant excitatory, inhibitory, or no change in activity to the peanut swab. (**G**) Proportion of excitatory and inhibitory responsive cells when the TMT swab was in the Near position (compared to Far position). (**H**) Correlation between Ca^2+^ activity dynamics of single *Pnoc*^BNST^ neurons and pupil size when mice were exposed to the TMT swab. (**I**) Correlation between Ca^2+^ activity dynamics of single *Pnoc*^BNST^ neurons and pupil size when mice were exposed to the Control swab (excited and inhibited as defined by their response to the TMT swab). (**J**) Proportion of excitatory and inhibitory responsive cells when the peanut swab was in the Near position (compared to Far position). (**K**) Correlation between Ca^2+^ activity dynamics of single *Pnoc*^BNST^ neurons and pupil size when mice were exposed to the peanut swab. Data shown as mean ± SEM. \*\*\*\**p<0.0001*.

Consistent with the valence of each odorant, the peanut oil swab produced a moderate increase in movement velocity when the odor was near, whereas the TMT swab produced an initial decrease in velocity (Figures S2A-B). Similar to pupil diameter, velocity and neural activity of *Pnoc*^BNST^ neurons showed significant correlations in neurons that showed a significant change in response (either excitation or inhibition) to mice exposed to the peanut swab (Figures S2C-E). The increase in correlation was very modest to TMT exposure and occurred only in neurons excited by the TMT swab, as compared to the control swab. In summary, we find that a large proportion of *Pnoc*^BNST^ neurons are correlated with measurements of arousal states, and the observed heterogeneity in response dynamics suggest that these neurons are composed of functionally distinct subtypes.

### *Pnoc*^BNST^ neurons drive arousal responses

Since we observed that excitatory responses were predominant to both aversive and rewarding odors, we next tested if optogenetic photoactivation of *Pnoc*^BNST^ neurons (Figure 4A-C) can induce a motivational state. We first evaluated if viral tools can activate *Pnoc*^BNST^ neuronal activity in a proficient and reliable manner. Whole-cell patch-clamp electrophysiological recordings in adBNST within ChR2-expressing *Pnoc*^BNST^ neurons (Figure 4B) showed that we could reliably photoactivate *Pnoc*^BNST^ neurons at 20 Hz with 100% spike fidelity (Figure 4C). To test if photoactivation of *Pnoc*^BNST^ neurons induced a place preference or aversion, freely moving mice were placed in a two-chambered arena to assess time spent in a chamber paired with photoactivation of *Pnoc*^BNST^ neurons (real-time place preference assay). Photoactivation of *Pnoc*^BNST^ neurons did not induce place aversion or place preference (Figure 4D-E), indicating that *Pnoc*^BNST^ neurons may not inherently drive approach/avoidance behaviors. Furthermore, photoactivation of *Pnoc*^BNST^ neurons did not induce changes in locomotion, as measured by velocity and freezing behavior (Figure S3A-C).

**Figure 4.**
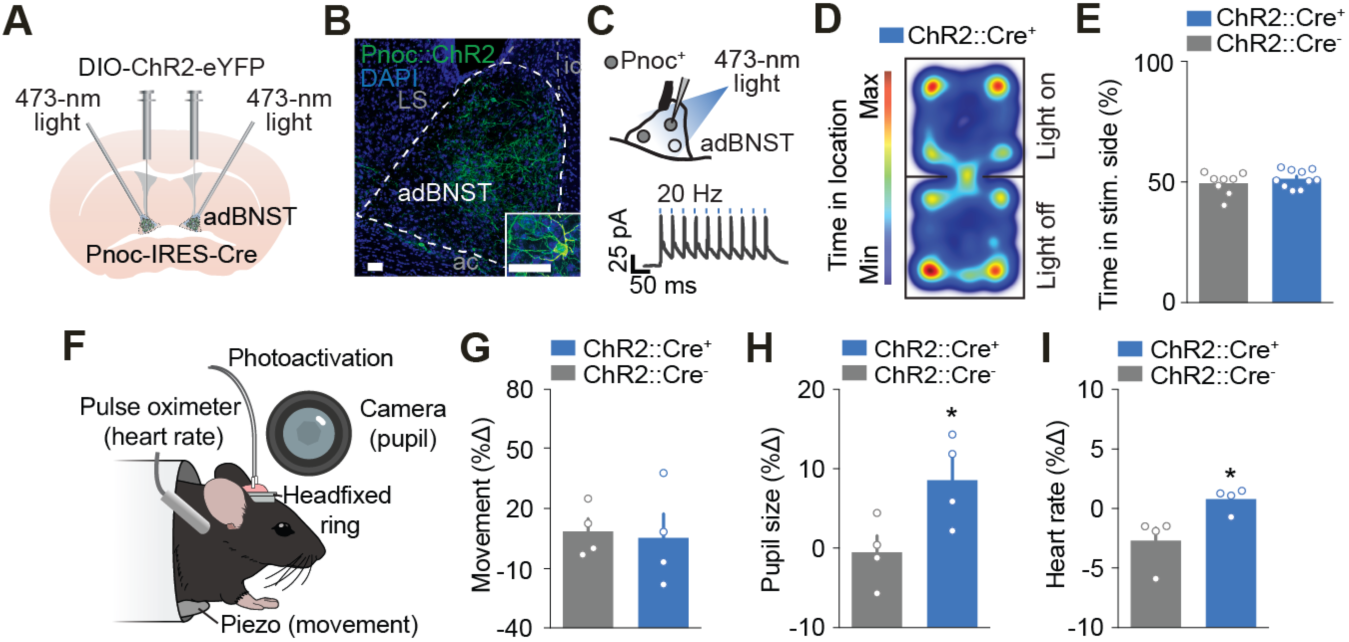
(**A**) Schematic of an injection of AAV5-EF1α-DIO-hChR2(H134R)-eYFP and implantation of fibers into the BNST of a *Pnoc*-IRES-*Cre* mouse. (**B**) Confocal image depicting expression of ChR2-eYFP in *Pnoc*^BNST^ neurons. Abbreviations: LS = lateral septum; ic = internal capsule; adBNST = anterodorsal BNST; ac = anterior capsule. *Inset:* Confocal image at high magnification depicting expression of ChR2-eYFP in *Pnoc*^BNST^ neurons. (**C**) Schematic of patch-clamp electrophysiology of ChR2-exressing *Pnoc*^+^ neurons (left). Sample neural response of a *Pnoc*^BNST^ neuron expressing ChR2 in response to blue light at 20 Hz (right). Group data showed 100% spike fidelity. (**D**) Sample heat map illustrating the location of a mouse during photoactivation of *Pnoc*^BNST^ neurons in the RTPP. (**E**) Group average for time in stimulation side during RTPP with photoactivation of *Pnoc*^BNST^ neurons. (**F**) Schematic of a head-fixed mouse in a cylindrical enclosure with an optical patch cable (photoactivation), a heart rate monitor (pulse oximeter), a movement monitor (piezo sensor), and a camera (pupil). (**G-I**) Group average for the change in movement (**G**), heart rate (**H**), and pupil size (**I**) with photoactivation of *Pnoc*^BNST^ neurons. Data shown as mean ± SEM. \**p<0.05*.

Therefore, we next wanted to test if photoactivation of *Pnoc*^BNST^ neurons was sufficient to increase physiological arousal. To accomplish this, we developed a stationary head-fixed preparation that allowed for the simultaneous measure of arousal responses in combination with optogenetics by transducing Cre-dependent channelrhodopsin into adBNST of *Pnoc*-Cre mice (Figure 4F). Photoactivation of *Pnoc*^BNST^ neurons did not alter movement, as measured by a piezoelectric sensor (Figure 4G), but significantly increased both pupil area (Figure 4H) and heart rate (Figure 4I). Photoactivation of these neurons also did not affect licking for a sucrose reward (Figure S3D-E), indicating that increasing arousal did not increase reward-seeking. Taken together, these data suggest that the activity of *Pnoc*^BNST^ neurons increases physiological arousal responses.

### *Pnoc*^BNST^ neurons exhibit diversity in both connectivity and genetic identity

Since we found heterogeneity in response dynamics with animals exposed to both arousal-inducing aversive and rewarding odors, we hypothesized that this heterogeneity may be due to diversity of connectivity and gene expression patterns within the *Pnoc*^BNST^ neuronal population. To investigate connectivity, we injected a Cre-dependent virus to express both a cytosolic and a presynaptic marker into adBNST. We found presynaptic labeling from *Pnoc*^BNST^ neurons within multiple compartments of BNST (evidenced by synaptophysin-mRuby expression), suggesting that these cells form local connections among various BNST subnuclei. Notably, we observed that presynaptic *Pnoc*^BNST^ terminals overlapped with both mGFP-labeled and unlabeled cells within the BNST, indicating that *Pnoc*^BNST^ neurons may form monosynaptic connections with both *Pnoc^+^* and *Pnoc^-^* neurons (Figure 5A). Whole-cell patch-clamp electrophysiological recordings revealed that light evoked inhibitory postsynaptic currents were detected in adBNST neurons following photostimulation *Pnoc*^BNST^ neurons, which was blocked by bath application of a GABA_A_ receptor antagonist (gabazine; Figure 5B), confirming local connectivity and the GABAergic phenotype of these cells. Furthermore, local inhibition arising from *Pnoc*^BNST^ activation was detected in a greater proportion of recorded eYFP*^-^* neurons (59%, putative non-*Pnoc*^BNST^ neurons), but still present in eYFP*^+^* neurons (31%, *Pnoc*^BNST^ neurons) (Figure 5C). Taken together, these data demonstrate that *Pnoc*^BNST^ neurons form local monosynaptic inhibitory connections with both putative *Pnoc*^-^ and *Pnoc*^+^ BNST neurons.

**Figure 5.**
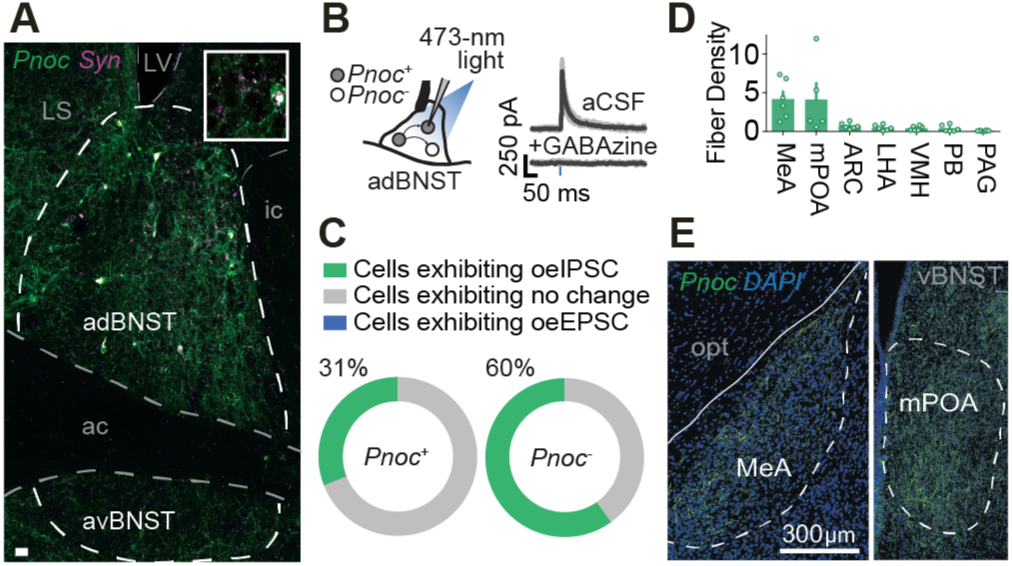
(**A**) Confocal image depicting the distribution of *Pnoc*^BNST^ somata and terminals (Syn = synaptophysin-mRuby) after local injection of AAVdj-HSyn-Flex-mGFP-2A-synaptophysin-mRuby. Inset shows higher magnification of adBNST. Abbreviations: LV = lateral ventricle; LS = lateral septum; adBNST = anterodorsal BNST; avBNST = anteroventral BNST; ic = internal capsule; ac = anterior commissure. (**B**) Schematic of patch-clamp electrophysiology of ChR2-exressing *Pnoc*^+^ neurons (left). Voltage-clamp traces from *Pnoc*^BNST^ neurons showing optically evoked inhibitory postsynaptic currents (oeIPSC) being blocked by GABA_A_ receptor antagonist GABAzine (right). Abbreviation: aCSF = artificial cerebral spinal fluid. (**C**) Proportion of *Pnoc^-^* and *Pnoc^+^* neurons exhibiting oeIPSC, no change, and optically evoked excitatory postsynaptic currents (oeEPSC). (**D**) Quantification of fiber density (% of area) across distal regions showing fiber labeling in animals expressing ChR2-EYFP in *Pnoc*^BNST^ neurons. Abbreviations: MeA = medial amygdala; mPOA = medial preoptic area; ARC = arcuate nucleus; LHA = lateral hypothalamic area; VMH = ventromedial hypothalamus; PB = parabrachial nucleus; PAG = periaqueductal grey. (**E**) Confocal image depicting fibers from animals expressing EYFP in *Pnoc*^BNST^ neurons in posterodorsal portion of medial amygdala at −1.94 mm from bregma (left) and medial preoptic area at −0.10 mm from bregma (right). Abbreviations: opt = optic tract; vBNST = ventral BNST.

To identify projection targets from *Pnoc*^BNST^ neurons, we labeled these neurons (including their axonal projections) and assessed the expression of their fluorescent markers in distal target regions. Distal axonal labeling was observed predominantly within the medial amygdala (MeA) and medial preoptic area (mPOA), with sparse to near absent labeling in other adBNST output regions including the arcuate nucleus (ARC), lateral hypothalamic area (LHA), ventromedial hypothalamus (VMH), parabrachial nucleus (PB), and periaqueductal grey (PAG) (Figure 5D-E) (Calhoon and Tye, 2015; Jennings et al., 2013a; Kim et al., 2013; Lebow and Chen, 2016). The MeA and mPOA are two regions critical for social motivation (Li et al., 2017; McHenry et al., 2017), therefore perhaps *Pnoc*^BNST^ projections to these regions may be involved in social arousal.

To address if *Pnoc*^BNST^ neurons are composed of distinct subpopulations of genetically identifiable neurons, we employed a single-cell sequencing approach using a droplet-based method (Drop-seq) (Macosko et al., 2015) that allowed us to capture mRNAs from 2,492 individual cells within BNST (median of 1435 genes/cell and 2257/cell unique transcripts; Figure 6A **&** S4A-F). We partitioned these cells into distinct clusters using cluster analysis based on gene expression patterns (Figure 6B **&** S4G). 11 out of 19 defined clusters expressed the canonical neuronal gene *Camk2b*, whereas remaining (8) clusters expressed known markers for non-neuronal cell types that defined astrocytes, oligodendrocytes, and oligodendrocyte precursor cells. Our single-cell sequencing approach revealed that *Vgat* is expressed more abundantly than *Vglut2* across all BNST neuronal clusters (Figure S4J). We found that 88% of *Pnoc*^BNST^ neurons were distributed among 4 of the 11 neuronal clusters (Figure 6C-D **&** S4H-I) that were differentiated by expression of somatostatin (*Som*), protein kinase C δ (*Pkcδ*), cholecystokinin (*Cck*), and the zic family member 1 (*Zic1*). Furthermore, little to no overlap (< 5%) was observed between *Pnoc*^BNST^ neurons and neuronal clusters defined by expression of forkhead box protein P2 (*Foxp2*), preproenkephalin (*Penk*), preprodynorphin (*Pdyn*), calbindin 2 (*Calb2*), corticotropin-releasing hormone (*Crh*), neurotensin (*Nts*), and Vglut3 (*Slc17a8*). FISH experiments corroborated a subset of our sequencing data (Figure 6E-F). In summary, these data suggest that *Pnoc*^BNST^ neurons can be further subdivided into at least 4 unique cell types identified by the coexpression of *Som, Pkcδ, Cck and Zic1*.

**Figure 6.**
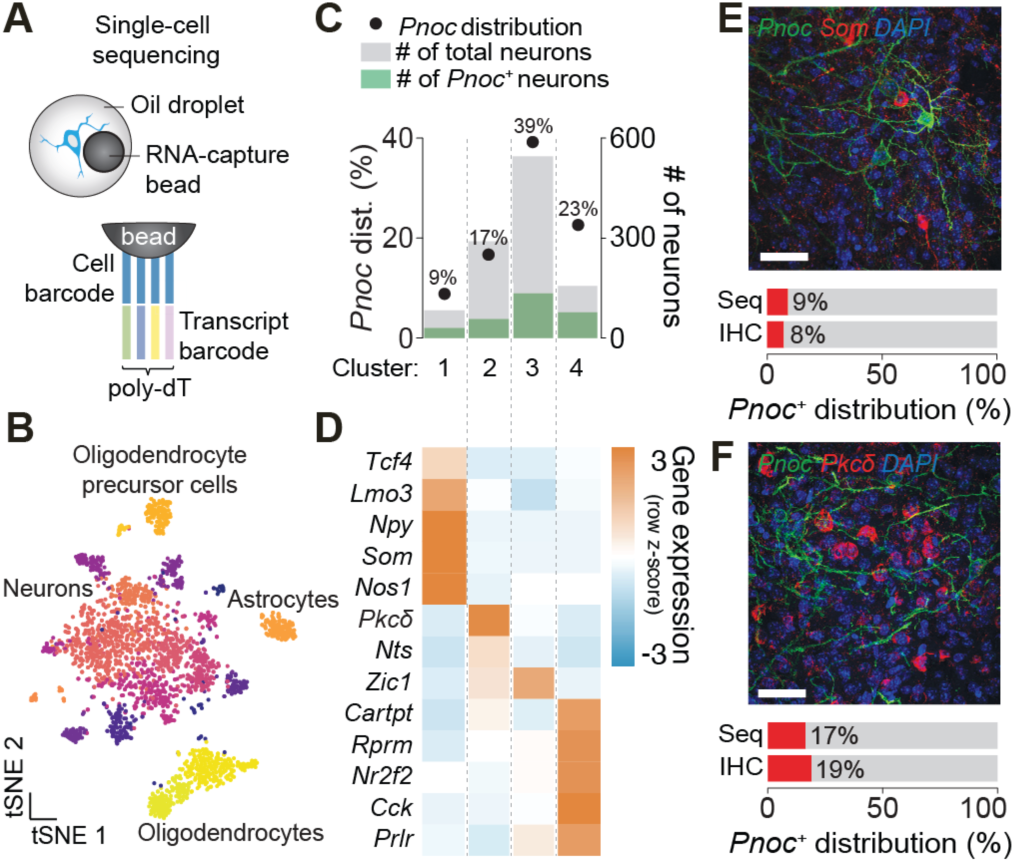
(**A**) Schematic of the droplet-based method (Drop-seq) used to sequence RNA from thousands of individual cells within the BNST. (**B**) Gene expression pattern of BNST visualized in tSNE space. Colors represent neuronal clusters. (**C**) Distribution of *Pnoc*^+^ neurons across clusters expressing >5% of *Pnoc*^+^ cells (left axis). Distribution of the number of total cells and *Pnoc*^+^ cells across the same clusters (right axis). (**D**) Heat map depicting expression of candidate marker genes for the same neuronal cluster in Figure 4F. (**E**) Confocal image depicting the overlap between the expression of *Pnoc* and *Som* within BNST neurons using immunohistochemistry (top). Distribution of *Som^+^* neurons quantified using either Drop-seq (Seq) or immunohistochemistry (IHC) (bottom). (**F**) Confocal image depicting the overlap between the expression of *Pnoc* and *Pkcδ* within BNST neurons using immunohistochemistry (top). Distribution of *Pkcδ^+^* neurons quantified using either Drop-seq (Seq) or immunohistochemistry (IHC) (bottom).

## DISCUSSION

In the present study, we found that *Pnoc* expression defines a subpopulation of GABAergic neurons within the BNST that are associated with changes in physiological arousal. *Pnoc*^BNST^ neurons encode the rapid increase in arousal that occur upon presentation of salient odorants that elicit both approach or avoidance behavior. Further, these neurons form local monosynaptic inhibitory connections with both *Pnoc*^-^ and *Pnoc*^+^ neurons within adBNST and also project to both MeA and mPOA. In agreement with the observed heterogeneity in *Pnoc*^BNST^ response dynamics, we found that *Pnoc*^BNST^ neurons can be divided into at least 4 genetically unique cell types that can be identified by co-expression of *Pnoc* with either *Som*, *Pkcδ*, *Cck* or *Zic1*. Taken together, these data show that *Pnoc*^BNST^ neurons have a critical role in driving rapid arousal responses that are characteristic of a variety of motivational states, and highlight the need for future studies to further unravel the heterogeneity within this genetically-identified neuron population.

Elevated anxiety is a maladaptive state that is associated with many neuropsychiatric conditions (Calhoon and Tye, 2015; LeDoux and Pine, 2016; Perusini and Fanselow, 2015). The manifestation of anxiety-like states includes both behavioral and physiological responses that need to occur rapidly in order to guide actions necessary for survival. Past research has developed an expansive literature on the neural circuits governing anxiety-like behavioral actions (for reviews see Calhoon and Tye, 2015; Harris and Gordon, 2015; Shin and Liberzon, 2010; Tovote et al., 2015), yet physiological arousal has received less attention. Excessive arousal responses as measured by increases in pupil size during threat exposure is commonly observed in patients suffering from anxiety disorders (Cascardi et al., 2015; Price et al., 2013). The presentation of negative emotional arousing images increases both pupil size and amygdala activity (as measured by BOLD signaling) (Hermans et al., 2013), but the relationship between these two variables has remained elusive. fMRI lacks both the temporal resolution needed to match the rapid changes in pupil size and the spatial resolution to identify subregions and more importantly individual neurons. In this study, we used calcium imaging to show that changes in the activity dynamics of individual *Pnoc*^BNST^ neurons are directly correlated with changes in pupil size, suggesting that these neurons may be a critical component for orchestrating excessive physiological arousal responses in pathological anxiety. Furthermore, our findings highlight a diversity of response dynamics and genetically identifiable subtypes within the *Pnoc*^BNST^ neuronal population that warrant further dissection. Experiments aimed at inhibiting or activating *Pnoc*^BNST^ subtypes during specific time windows following presentation of arousal-inducing stimuli will aid in understanding the role of this ensemble in orchestrating rapid arousal responses.

We also found that *Pnoc*^BNST^ neurons consist of an interconnected microcircuit of GABAergic neurons within the BNST that may be classified by the expression of distinct genetic markers (*Som*, *Prkcd*, *Cck*, and *Zic1*). This further suggests that either functionally distinct subtypes of *Pnoc*^BNST^ neurons exist or molecularly distinct subtypes of BNST neurons share a similar function. Therefore, future studies are needed to systematically assess the functional role of *Pnoc*^BNST^ neuronal subtypes and their role in rapid arousal responses. For example, co-expression of *Npy* and *Som* has been previously reported throughout the entire amygdala (McDonald, 1989), suggesting that both of these markers identify a single neuronal cell type. Our data identify a similar neuronal cluster characterized by the co-expression of *Npy* and *Som*. It was previously shown that *Npy*-expressing neurons have a specific projection output to the preoptic region of the hypothalamus (Pompolo et al., 2005). Our data show that at least a subset of *Pnoc*^BNST^ neurons share this projection. Taken together, perhaps these three genetic markers (*Pnoc*, *Npy*, *Som*) may be used to target the sub-population of *Pnoc*^BNST^ neurons that project from BNST to mPOA. Considering the role of mPOA in social avoidance/approach behavior (McHenry et al., 2017), this projection could be important for social arousal.

A recent study showed that local photoactivation of all *Som*^BNST^ neurons drives anxiety-mediated avoidance in the EPM (Ahrens et al., 2018). Therefore, investigating the role of how *Pnoc^+^*/*Som^+^* and *Pnoc^+^*/*Som*^-^ neurons might differ in the regulation of anxiety-like behavior and arousal responses deserves further attention. Additionally, future studies using intersectional genetic approaches to target *Pnoc* and either *PKCd*, *Cck*, or *Zic1* neurons could help to further characterize the heterogeneity of activity responses we observed in distinct *Pnoc*^BNST^ subsets. It is also equally important to delineate how *Pnoc*^BNST^ neurons may interact with other local *Pnoc^+^* neurons and local *Pnoc^-^* neuron clusters, such as *Foxp2*^BNST^, *Penk*^BNST^, *Pdyn*^BNST^, *Calb2*^BNST^, *Crh*^BNST^*, Nts*^BNST^*, and Vglut3*^BNST^ (Gafford and Ressler, 2015; Hammack et al., 2009; Kash et al., 2015; Lebow and Chen, 2016; McElligott and Winder, 2009; Nguyen et al., 2016).

BNST neurons have also been distinguished by their projection targets in previous studies. For instance, PBN-projecting neurons regulate autonomic arousal states as measured by respiration, whereas neurons projecting to the LHA regulate anxiety-like behavior in the EPM (Kim et al., 2013; Kodani et al., 2017). A recent study showed that BNST neurons that project to the LHA can be further subdivided by the expression of the neuropeptidergic genes corticotropin-releasing hormone (*Crh*^BNSTàLHA^) and cholecystokinin (*Cck*^BNSTàLHA^). These neurons show an increase in average calcium activity specific to a rewarding (female mouse urine) or aversive odorant (TMT), respectively (Giardino et al., 2018). Furthermore, chemogenetic activation of *Vgat* expressing neurons within the BNST increases anxiety-like behavior and leads to activation of the locus coeruleus (LC) (Mazzone et al., 2018). Although these phenotypes are similar to our findings with *Pnoc*^BNST^ neurons, we did not observe appreciable projections from *Pnoc*^BNST^ neurons to either PBN, LHA or LC, indicating that *Pnoc*^BNST^ neurons may be distinct from both PBN-, LHA- and LC-projecting neurons. Nonetheless, whether local interactions between *Pnoc*^BNST^ neurons and either PBN-, LHA-, or LC-projecting neurons within BNST exist remains an open question that warrants further investigation.

Using advanced tools to probe neurons with single-cell resolution, we discovered that *Pnoc*^BNST^ neurons encode rapid changes in physiological arousal responses. However, these neurons are likely only a piece of the complex mosaic of cell types within BNST that contribute towards arousal responding and motivational states. Further investigations into how *Pnoc*^BNST^ neurons and other BNST cell types differentially and synergistically control rapid arousal responses will shed light onto how BNST and the larger network of brain regions that regulate motivational states contribute to the development and perpetuation of neuropsychiatric disorders characterized by maladaptive motivational states.

---

**Figure S1.**
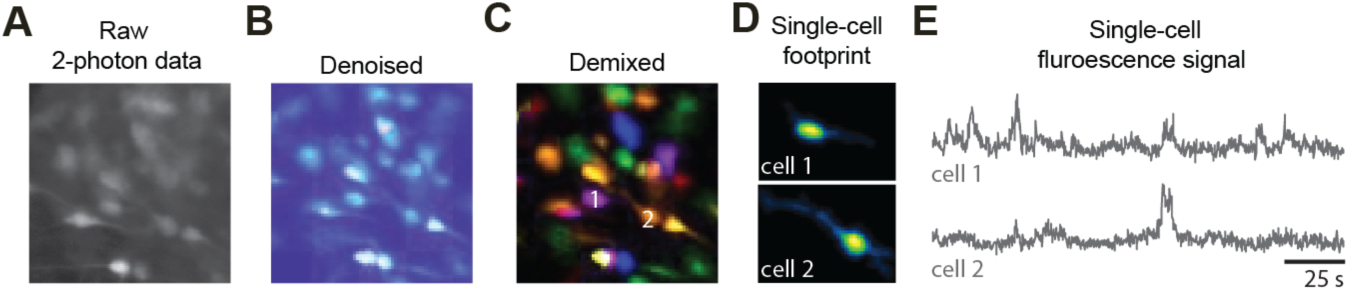
(**A-E**) Analysis pipeline of calcium imaging data using constrained nonnegative matrix factorization (CNMF) for extracting single-cell fluorescence signals from imaging data (Zhou et al., 2018).

**Figure S2.**
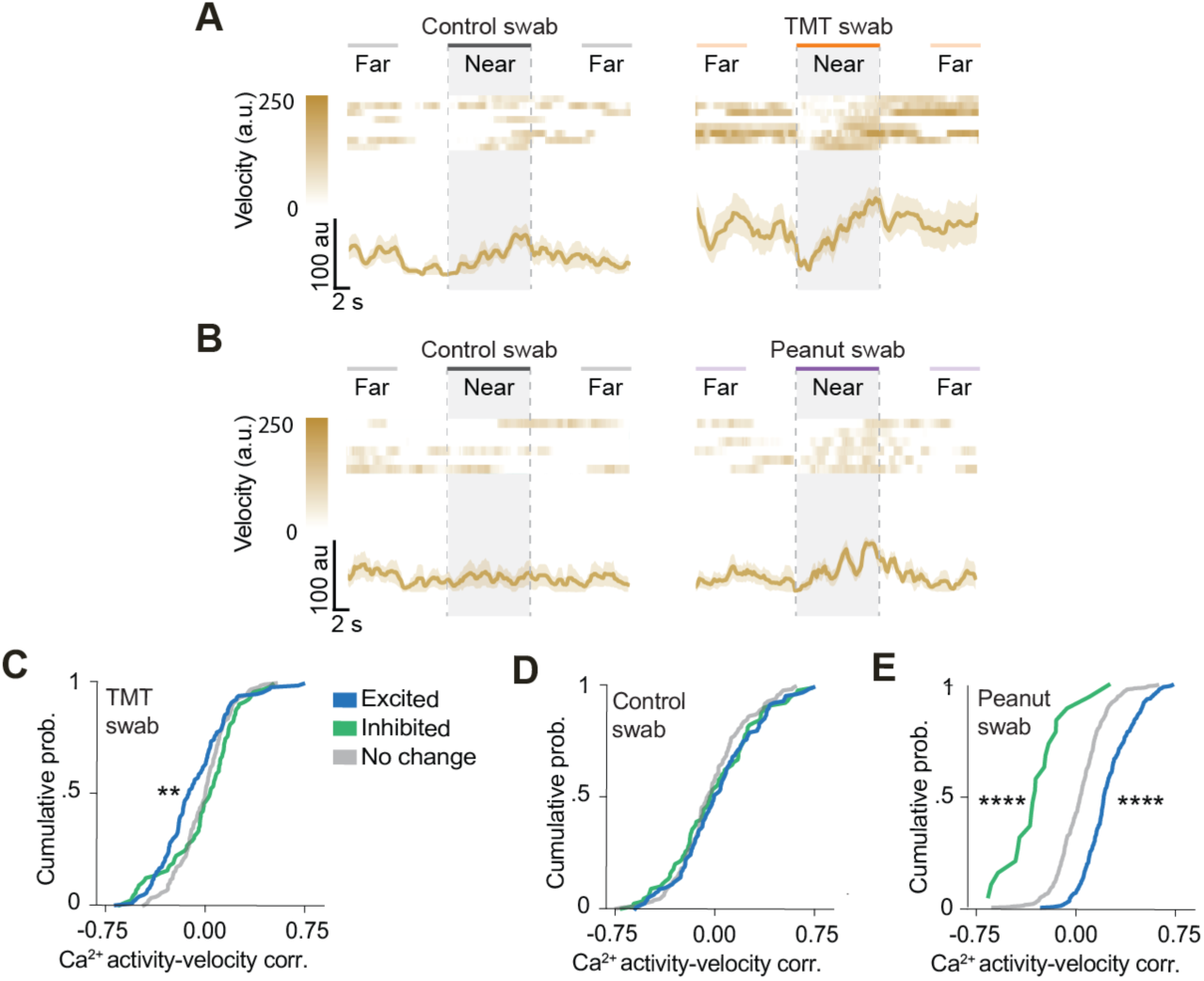
(**A**) Heat map of individual data (top) and average group data (bottom) for velocity responses to the control and TMT swabs. (**B**) Heat map of individual data (top) and average group data (bottom) for velocity responses to the control and peanut swabs. (**C**) Correlation between Ca^2+^ activity dynamics of single *Pnoc*^BNST^ neurons and velocity when mice were exposed to the TMT swab. (**D**) Correlation between Ca^2+^ activity dynamics of single *Pnoc*^BNST^ neurons and velocity when mice were exposed to the control swab (excited and inhibited as defined by their response to the TMT swab). (**E**) Correlation between Ca^2+^ activity dynamics of single *Pnoc*^BNST^ neurons and velocity when mice were exposed to the peanut swab. Data shown as mean ± SEM. \*\**p<0.01,* \*\*\*\**p<0.0001*.

**Figure S3.**
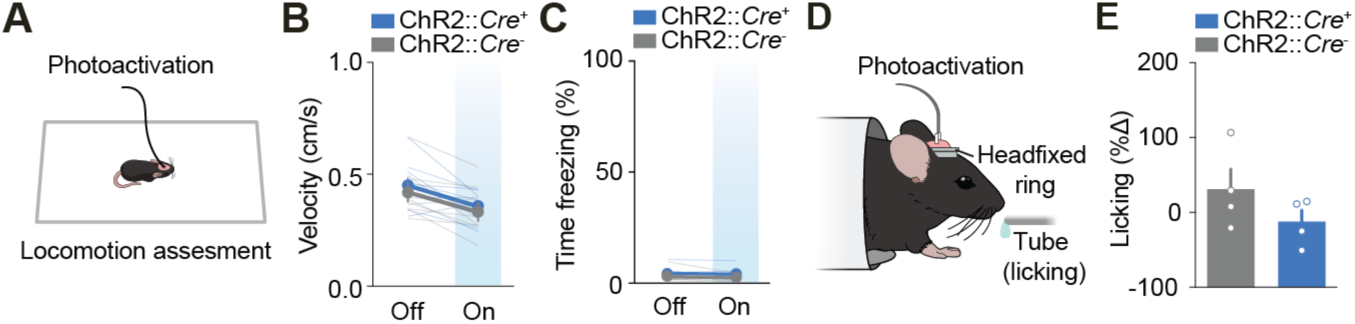
(**A**) Schematic of a tethered (for optogenetics) freely moving mouse in a small rectangular arena to assess locomotion. (**B**) Group average for velocity with photoactivation of *Pnoc*^BNST^ neurons. (**C**) Group average for time freezing with photoactivation of *Pnoc*^BNST^ neurons. (**D**) Schematic of a head-fixed mouse in a cylindrical enclosure with an optical patch cable (optogenetics) and a tube (licking) for sucrose delivery. (**E**) Group average for the change in licking with photoactivation of *Pnoc*^BNST^ neurons. Data shown as mean ± SEM.

**Figure S4.**
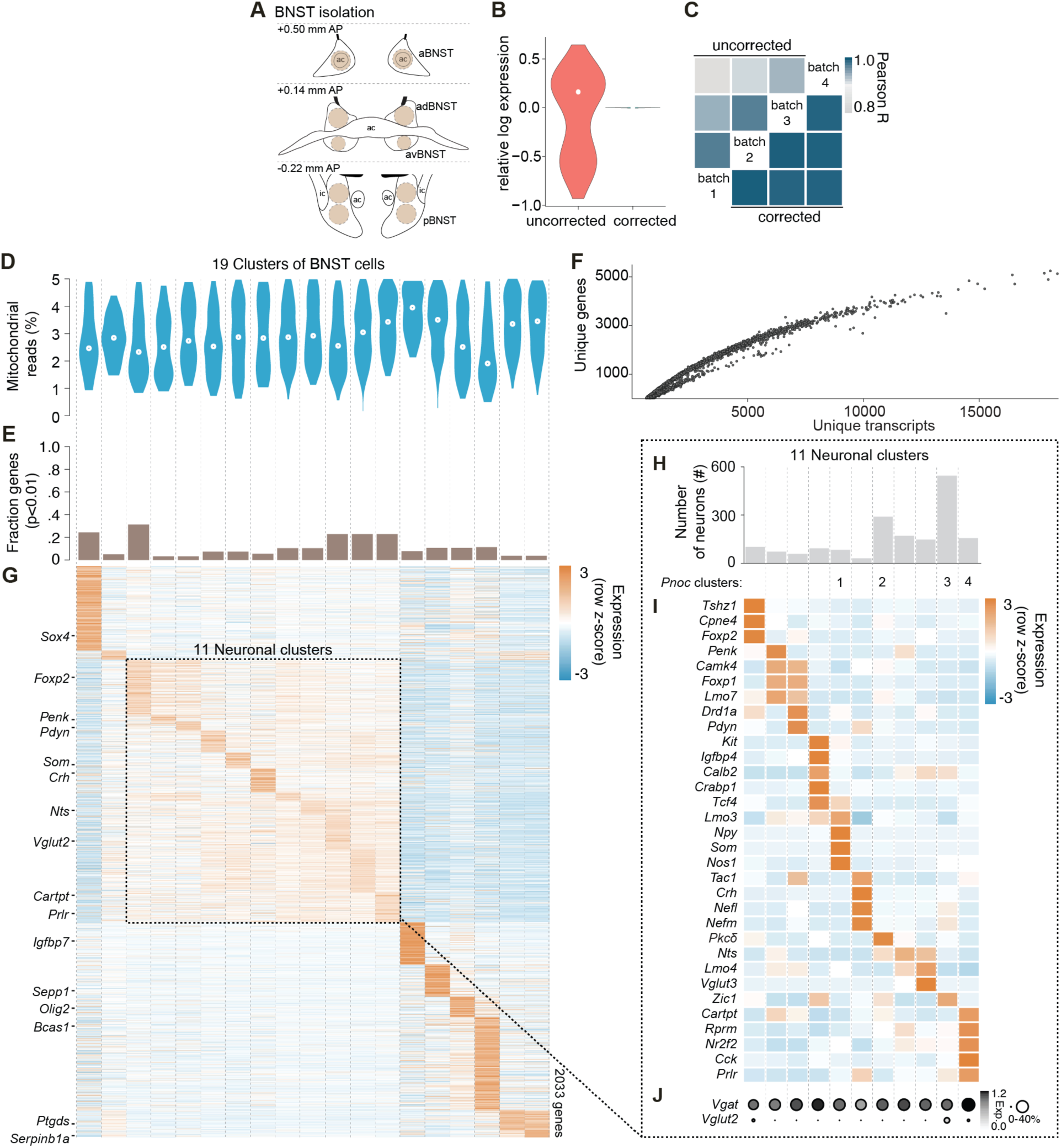
(**A**) Schematic of tissue isolated from BNST. (**B**) Relative log expression normalization across cells (**C**) and mean gene expression correlations across batches following parametric batch correction. (**D**) Percent mitochondrial reads distribution across clusters. (**E**) Fraction of significant genes marking each cluster as determined using a likelihood-ratio test for single-cell data. (**F**) Individual cells plotted by the number of unique genes and unique transcripts detected. (Median = 1434.5 unique genes, 2257 unique transcripts). (**G**) Gene mean expression across all cell clusters. The 11 neuronal clusters are highlighted. (**H**) Distribution of all cells across clusters. (**I**) Heat map depicting expression of candidate marker genes for each neuronal cluster. (**J**) Expression of *Vgat* and *Vglut2* across all neuronal clusters. Color represents normalized gene expression level. Size corresponds to proportion of neurons that expressed gene.

## SUPPLEMENTAL INFORMATION

Supplemental information includes 4 figures and one table and can be found with this article online.

## AUTHOR CONTRIBUTIONS

JRR, RLU, and GDS designed the experiments.

JRR, MLB, JMO, HN, JER, XZ, and OK conducted the experiments.

HTC, TLK, and MRB provided either critical reagents and/or feedback.

RLU, JRR, MLB, JMO, VMKN, HN, JAM, and GDS analyzed the data.

JRR, RLU, and GDS wrote the manuscript with comments from all co-authors.

## ACKNOWLEDGEMENTS

We thank Hiroyuki K. Kato, Anthony Burgos-Robles, Maria M. Diehl, Fabricio H. Do-Monte, Ivan Trujillo-Pisanty and Gregory J. Quirk for helpful discussions and comments on the manuscript. We thank K. Deisseroth and the GENIE project at Janelia Research Campus for viral constructs. This work was supported by grants from the National Institute of Mental Health (F32-MH113327, J.R.R.; F30-MH115693, R.L.U.; T32-MH093315 & K99-MH115165, J.A.M.), National Institute of Neurological Disorders and Stroke (T32-NS007431, R.L.U.), National Institute of Drug Abuse (F32-DA041184, J.M.O., R37-DA032750 & R01-DA038168, G.D.S.), Children’s Tumor Foundation (016-01-006, J.E.R.), Brain and Behavior Research Foundation (G.D.S.), Yang Biomedical Scholars Award (G.D.S.), Foundation of Hope (G.D.S.), UNC Neuroscience Center (G.D.S.; Helen Lyng White Fellowship, V.M.K.N.), UNC Neuroscience Center Microscopy Core (P30-NS045892) and UNC Department of Psychiatry (G.D.S.).

## KEY RESOURCES TABLE

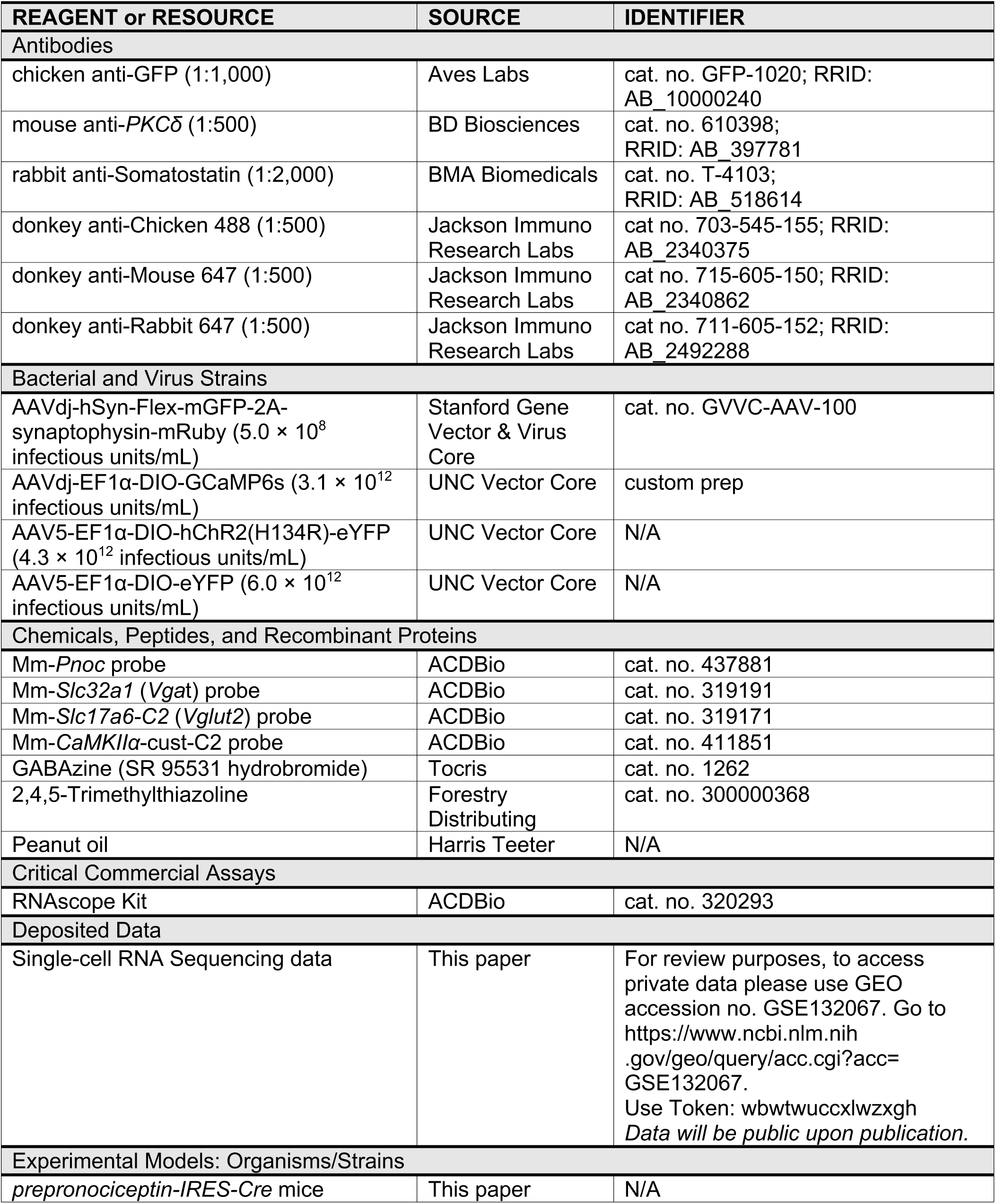
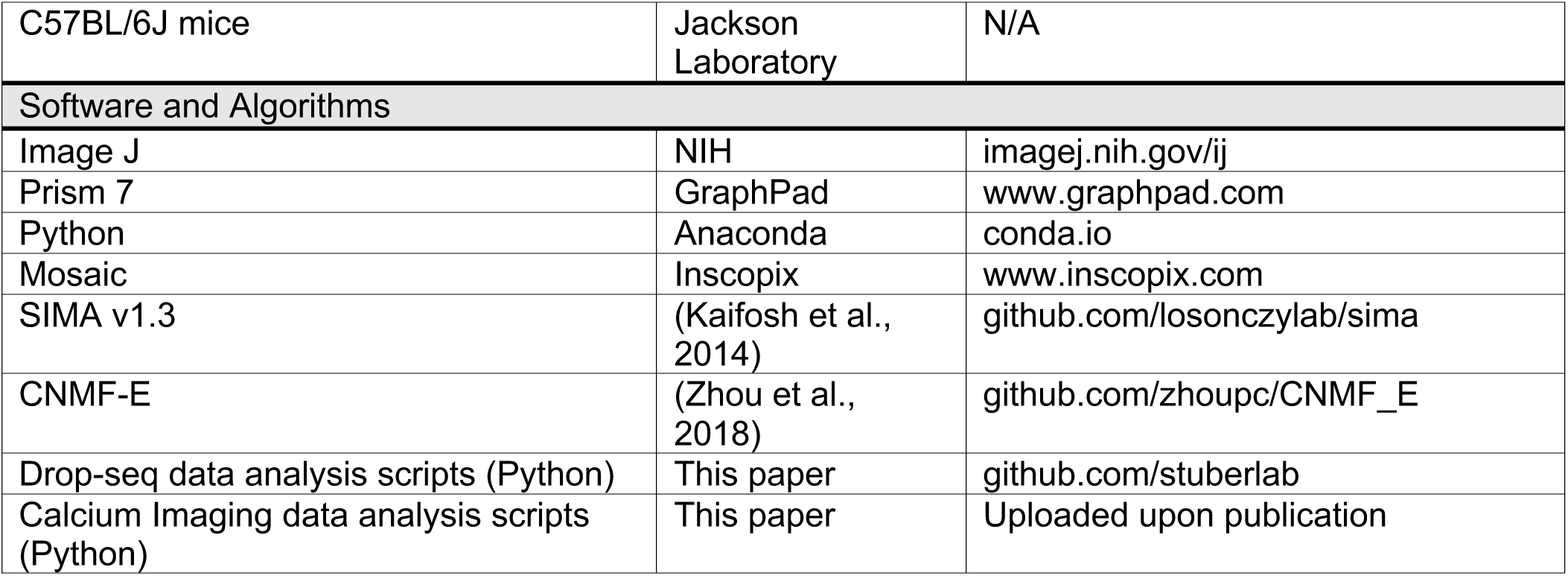

TABLE S1.

| **STATISTICAL ANALYSIS** |  |  |  |  |
| --- | --- | --- | --- | --- |
| **Figure** | **# of Samples** | **Test** | **Test statistic** | **p value** |
| **Figure 1** |  |  |  |  |
| K; Evoked fluorescent/spike relationship | n = 10 *Pnoc*<sup>+</sup> neurons | Pearson correlation coefficient | r = 0.996 | *p*<0.001 |
| **Figure 2** |  |  |  |  |
| B; Time in odor zone when exposed to the control and TMT swabs | n = 4 wild type mice | Paired t-test | *t*<sub>3</sub>=6.518 | *p*=0.0073 |
| C; Time in odor zone when exposed to the control and peanut swabs | n = 4 wild type mice | Paired t-test | *t*<sub>3</sub>=6.277 | *p*=0.0082 |
| E; Pupil size when exposed to the control and TMT swabs | n = 4 wild type mice | Paired t-test | *t*<sub>3</sub>=3.604 | *p*=0.0367 |
| F; Pupil size when exposed to the control and peanut swabs | n = 4 wild type mice | Paired t-test | *t*<sub>3</sub>=4.298 | *p*=0.0232 |
| **Figure 3** |  |  |  |  |
| H; Correlation between Ca<sup>2+</sup> activity and pupil size when mice were exposed to the TMT swab (responses defined by exposure to the TMT swab) | n = 401 *Pnoc*<sup>+</sup> neurons | Mann-Whitney U test, significance level adjusted using Bonferroni correction | Excited vs No Change: *U*=2527.0<br>Inhibited vs No Change: *U*=1671.0 | *p*=6.5808e-31<br>*p*=1.0754e-25 |
| I; Correlation between Ca<sup>2+</sup> activity and pupil size when mice were exposed to the control Swab (responses defined by exposure to the TMT swab) | n = 401 *Pnoc*<sup>+</sup> neurons | Mann-Whitney U test, significance level adjusted using Bonferroni correction | Excited vs No Change: *U*=11474.0<br>Inhibited vs No Change: *U*=7683.0 | *p*=2.9406<br>*p*=1.6152 |
| K; Correlation between Ca<sup>2+</sup> activity and pupil size when mice were exposed to the peanut swab (responses defined by exposure to the peanut swab) | n = 779 *Pnoc*<sup>+</sup> neurons | Mann-Whitney U test, significance level adjusted using | Excited vs No Change: *U*=24140.0<br>Inhibited vs No Change: *U*=971.0 | *p*=8.8104e-56<br>*p*=8.9200e-8 |
|  |  | Bonferroni correction |  |  |
| Figure 4 |  |  |  |  |
| E; Time in stimulation side during real time place preference | n = 8 ChR2-Cre<sup>-</sup> mice, n = 10 ChR2-Cre<sup>+</sup> mice | Two sample unpaired t-test | $t_{16}=0.9711$ | $p=0.3459$ |
| G; Movement | n = 4 ChR2-Cre<sup>-</sup> mice, n = 4 ChR2-Cre<sup>+</sup> mice | Two sample unpaired t-test | $t_6=0.240$ | $p=0.8185$ |
| H; Pupil size | n = 4 ChR2-Cre<sup>-</sup> mice, n = 4 ChR2-Cre<sup>+</sup> mice | Two sample unpaired t-test | $t_6=2.616$ | $p=0.0398$ |
| I; Heart rate | n = 4 ChR2-Cre<sup>-</sup> mice, n = 4 ChR2-Cre<sup>+</sup> mice | Two sample unpaired t-test | $t_6=2.928$ | $p=0.0264$ |
| Figure 5 |  |  |  |  |
| B; GABA<sub>A</sub> block | n = 4 *Pnoc*<sup>+</sup> neurons | Paired t-test | $t_3=4.83$ | $p=0.0064$ |
| Supplementary Figure 2 |  |  |  |  |
| C; Correlation between Ca<sup>2+</sup> activity and velocity when mice were exposed to the TMT swab (responses defined by exposure to the TMT swab) | n = 401 *Pnoc*<sup>+</sup> neurons | Mann-Whitney U test, significance level adjusted using Bonferroni correction | Excited vs No Change: $U=8790.0$<br>Inhibited vs No Change: $U=7438.0$ | $p=0.0013$<br>$p=0.9216$ |
| D; Correlation between Ca<sup>2+</sup> activity and velocity when mice were exposed to the control Swab (responses defined by exposure to the TMT swab) | n = 401 *Pnoc*<sup>+</sup> neurons | Mann-Whitney U test, significance level adjusted using Bonferroni correction | Excited vs No Change: $U=101550.0$<br>Inhibited vs No Change: $U=7818.0$ | $p=0.6720$<br>$p=2.0868$ |
| E; Correlation between Ca<sup>2+</sup> activity and velocity when mice were exposed to the peanut swab (responses defined by exposure to the peanut swab) | n = 779 *Pnoc*<sup>+</sup> neurons | Mann-Whitney U test, significance level adjusted using Bonferroni correction | Excited vs No Change: $U=28737.0$<br>Inhibited vs No Change: $U=995.0$ | $p=8.4642e-46$<br>$p=1.1500e-7$ |
| Supplementary Figure 3 |  |  |  |  |
| B; Velocity | n = 8 ChR2-Cre<sup>-</sup> mice, n = 9 ChR2-Cre<sup>+</sup> mice | Two-way ANOVA | Group: $F_{(1,15)}=0.3553$<br>Laser: | $p=0.5600$ |
| | | | $F_{(1,15)}=16.23$<br>*Interaction:*<br>$F_{(1,15)}=0.103$ | $p=0.0011$<br>$p=0.7527$ |
| | | Bonferroni | Laser Off<br>Laser On | $p=0.999$<br>$p=0.999$ |
| C; Time freezing | n = 9 ChR2-Cre<sup>-</sup> mice,<br>n = 8 ChR2-Cre<sup>+</sup> mice | Two-way<br>ANOVA | *Group:*<br>$F_{(1,15)}=2.123$<br>*Laser:*<br>$F_{(1,15)}=1.817$<br>*Interaction:*<br>$F_{(1,15)}=0.103$ | $p=0.1658$<br>$p=0.1977$<br>$p=0.7527$ |
| | | Bonferroni | Laser Off<br>Laser On | $p=0.4172$<br>$p=0.2962$ |
| E; Licking | n = 4 ChR2-Cre<sup>-</sup> mice,<br>n = 4 ChR2-Cre<sup>+</sup> mice | One sample<br>t-test | $t_6=1.376$ | $p=0.2179$ |

## STAR METHODS

### CONTACT FOR REAGENTS AND RESOURCE SHARING

Information and request for reagents may be directed and will be fulfilled by the corresponding author Garret D. Stuber.

### EXPERIMENTAL MODEL AND SUBJECT DETAILS

#### Animals

Adult (25-30g) male prepronociceptin*-*IRES*-Cre* (*Pnoc*-IRES-Cre) or wild type mice (C57 BL6/J) were independently housed and maintained on a reverse 12-hr light-dark cycle (lights off at 08:00 AM) with *ad libitum* access to food and water. Behavior was tested during the dark cycle. All procedures were conducted in accordance with the Guide for the Care and Use of Laboratory Animals, as adopted by the National Institute of Health, and with the approval of the Institutional Animal Care and Use Committee from the University of North Carolina at Chapel Hill.

### METHOD DETAILS

#### Fluorescent *In Situ* Hybridization

For processing tissue samples for *in situ* hybridization, mice were anesthetized with isoflurane (3.5-4.0%) vaporized in pure oxygen (1 L/min), rapidly decapitated and brains immediately extracted, and flash frozen on dry ice. 18 μm thick coronal sections were collected with a cryostat under RNase-free conditions, fixed in 4% PFA for 15 min at 4°C, dehydrated in serial concentrations of ethanol (50-100%), and processed according to instructions provided in the RNAscope kit (Advanced Cell Diagnostics, Newark, CA). Sections were hybridized with the following mixed probes: *Pnoc* (Mm-*Pnoc*, cat. no. 437881), *Vgat* (Mm-*Slc32a1*, cat. no. 319191), *Vglut2* (Mm-*Slc17a6*-C2, cat. no. 319171), *CaMKIIα* (Mm-*Camk2a*-cust-C2, cat. no. 411851). Hybridization probes used can also be found in supplementary information (Table S1). Following amplification, sections were counterstained with DAPI and cover slipped for subsequent confocal microscopy and counted using ImageJ software.

#### Immunohistochemistry

For processing tissue samples for immunohistochemistry, mice were euthanized with pentobarbital (50 mg.kg, 1.p.) and transcardially perfused with 0.01 M phosphate-buffered saline (PBS) and 4% paraformaldehyde (PFA). Tissue was fixed overnight in PFA at 4°C, cryoprotected with 30% sucrose in PBS, and 40 μm thick coronal sections were collected with a cryostat. Immunochemistry was performed in *Pnoc*-IRES-*Cre* mice using the following primary (kept overnight at 4 °C) and secondary (kept at room temperature for 2 h) antibodies: chicken-anti-GFP (1:1,000; Aves labs, Tigard, OR), donkey anti-chicken 488 (1:500; Jackson Immuno Research Labs, West Grove, PA), mouse anti-PKCδ (1:500; BD Biosciences, Fanklin Lakes, NJ), donkey anti-mouse 647 (1:500; Jackson Immuno Research Labs, West Grove, PA), rabbit anti-Somatostatin (1:2,000; BMA Biomedicals, Switzerland), and donkey anti-rabbit 647 (1:500; Jackson Immuno Research Labs, West Grove, PA). Antibodies used with dilutions can also be found in supplementary information (Table S1). Immunoprocessing procedures were done as previously described (Jennings et al., 2013a), and sections were counterstained with DAPI and cover slipped for subsequent confocal microscopy and counted using ImageJ software.

#### Confocal Microscopy

A confocal microscope (Zeiss LSM 780, Carl Zeiss, San Diego, CA) with either a 20x (air), 40x (air), or a 63x (oil) objective was used to capture images. Software (Zen Software, Carl Zeiss, Jena, Germany) settings were optimized for each experiment. In general, z-stacks were acquired in less than 1 μm increments and the maximum intensity projection of tiled images were used for representative images or for quantification purposes.

#### Viral Constructs

All viral constructs [Cre-inducible AAVdj-EF1α-DIO-GCaMP6s (3.1 x 10^12^ infectious units/mL), AAV5-EF1α-DIO-hChR2(H134R)-eYFP (4.3 x 10^12^ infectious units/mL), AAV5-EF1α-DIO-eYFP (6.0 x 10^12^ infectious units/mL), and AAVdj-hSyn-Flex-mGFP-2A-synaptophysin-mRuby (5.0 x 10^8^ infectious units/mL) were packaged by the UNC Vector Core and can also be found in supplementary information (Table S1).

#### Surgery and Histology

Mice were anesthetized with isoflurane (0.8-1.5%) vaporized in pure oxygen (1 l/min^-1^) and placed in a stereotaxic frame (David Kopf Instruments, Tujunga, CA). Ophthalmic ointment (Akorn, Lake Forest, IL) and topical anesthetic (2% lidocaine; Akorn, Lake Forest, IL) were applied during surgeries, and subcutaneous administration of saline (0.9% NaCL in water) were administered to prevent dehydration. Microinjections using injection needles (33 gauge) connected to a 2 uL syringe (Hamilton Company, Reno, NV) were used to deliver viruses into the anterior dorsal portion of the bed nucleus of the stria terminalis (adBNST; 500 nl per side; relative to bregma: +0.14 AP, +/-0.95 ML, DV −4.20 DV). For calcium imaging studies, unilateral virus injections were made into adBNST. To allow subsequent imaging of BNST neurons, a microendoscope [a gradient refractive index (GRIN) lens, 0.6 mm in diameter, 7.3 mm in length; Inscopix, Palo Alto, CA] was implanted 0.2 mm dorsal to the adBNST target site.

For optogenetic studies, bilateral virus injections were made into BNST, and an optical fiber was implanted with a 10° angle approximately 0.5 mm above the BNST. For experiments involving head-fixed behavior, a custom-made head-ring (stainless steel; 5 mm ID, 11 mm OD) was attached to the skull during surgery to allow head-fixation. Following surgeries, mice were given acetaminophen in their drinking water for 2 days and were allowed to recover with access to food and water *ad libitum* for at least 21 days. Following behavioral experiments, all cohorts were euthanized and perfused, tissue was extracted and 40 μm thick coronal sections collected with a cryostat, counterstained with DAPI and cover slipped for verification of viral expression and fiber/lens placement.

#### Patch-Clamp Electrophysiology

Mice were anesthetized with pentobarbital (50mg/kg) before transcardial perfusion with ice-cold sucrose cutting solution containing the following (in mM): 225 sucrose, 119 NaCl, 1.0 NaH_2_PO_4_, 4.9 MgCl_2_, 0.1 CaCl_2_, 26.2 NaHCO_3_, 1.25 glucose, 305 mOsm. Brains were then rapidly removed, and 300 μm thick coronal sections containing BNST were taken using a vibratome (Leica, VT 1200, Germany). Sections were then incubated in aCSF (32°C) containing the following (in mM): 119 NaCl, 2.5 KCl, 1.0 NaH_2_P0_4_, 1.3 MgCl, 2.5 CaCl_2_, 26.2 NaHCO_3_, 15 glucose, ∼306 mOsm. After an hour of recovery, slices were constantly perfused with aCSF (32°C) and visualized using differential interference contrast through a 40x water-immersion objective mounted on an upright microscope (Olympus BX51WI, Center Valley, PA). Recordings were obtained using borosilicate pipettes (3–5 ΜΩ) back-filled with internal solution containing the following (in mM): 130 K gluconate, 10 KCl, 10 HEPES, 10 EGTA, 2 MgCl_2_, 2 ATP, 0.2 GTP (pH 7.35, 270-285 mOsm.

Current-clamp recordings were obtained from GCaMP6s-expressing *Pnoc*^BNST^ neurons to identify how action potential frequency correlated with GCaMP6s fluorescence as previously described (Otis et al., 2017). Specifically, to determine how elevations in action potential frequency influence GCaMP6s fluorescence, a 1 second train of depolarizing pulses (2 nA, 2 ms) was applied at a frequency of 1, 2, 5, 10, or 20 Hz. During electrophysiological recordings, GCaMP6s fluorescence dynamics were visualized using a mercury lamp (Olympus U-RFL-T, Center Valley, PA) and a microscope-mounted camera (QImaging, optiMOS, Canada). Imaging data were acquired using Micro-Manager and extracted through hand-drawn ROIs for each recorded neuron using ImageJ. In addition to these experiments, we also performed current-clamp recordings to determine the spike fidelity of *Pnoc*^BNST^ ChR2-expressing neurons during optogenetic stimulation. To do so, neurons were held at resting membrane potential (n=7), and a blue LED (490nm; 1 mW) was presented in a series of 10 pulses (5 ms per pulse) at 1, 5, 10, and 20 Hz. We found that every pulse evoked an action potential for all neurons, suggesting 100% spike fidelity across cells.

Voltage-clamp recordings were obtained from BNST ChR2-expressing neurons (*Pnoc^+^*), and BNST non-ChR2 expressing neurons (*Pnoc^-^*) to identify local synaptic innervation of *Pnoc*^BNST^ neurons. To determine if a neuron was *Pnoc^+^*, we held all cells at −70mV and tested for the presence of ChR2 by displaying a blue LED (490nm; <1mW) for 1s. In the case that a long, stable inward current was evoked for the duration of that sweep, the neuron was confirmed to be *Pnoc^+^* and ChR2^+^ (n=26). Otherwise, the neuron was assumed to be *Pnoc^-^* and ChR2^-^ (n=37). We did not detect the presence of any transient, optically-evoked excitatory postsynaptic current (oeEPSC) during these sweeps, suggesting that *Pnoc*^BNST^ neurons do not release excitatory transmitters within this circuit. Next, we held all neurons at the reversal potential for ChR2 (+5 to +15 mV for *Pnoc^+^* neurons; +10 mV for *Pnoc^-^* neurons) and tested for the presence of an optically-evoked inhibitory postsynaptic current (oeIPSC) by displaying the blue LED for 5 ms. In a subset of cells, we tested whether the oeIPSC was mediated by GABA_A_ receptors by bath-applying GABAzine (10 uM) for 5 minutes. For all voltage-clamp experiments, data acquisition occurred at a 10 kHz sampling rate. All patch-clamp recordings were made through a MultiClamp 700B amplifier connected to a Digidata 1440A digitizer (Molecular Devices, San Jose, CA) and analyzed using Clampfit 10.3 (Molecular Devices, San Jose, CA).

#### Odor Preference in Freely-moving Mice

Mice were habituated to have a square block holder in their home cage for 2 days prior to testing. The day of testing, a cotton swab was placed in the square block holder located in an upright position 4 in from the home cage floor on one of the sides (sides were alternated across all mice). Mice behavior was recorded for a 5-min period after placing 2.5 μl of water (distilled H_2_O) in the cotton swab, followed by placing either 2.5 μl of TMT or 2.5 μl of Peanut oil (same as head-fixed experiment) in the cotton swab. Distance to odor (cm, max: 25 cm), time spent freezing (s), and velocity (cm/s) were calculated using automated tracking software (Ethovision XT 11, Noldus, Leesburg, VA). Similar to head-fixed odor exposure experiments, a low dose of TMT was used to minimize freezing responses and maintain ambulation.

Pupil recordings were made in freely moving animals using the same camera system used for head-fixed experiments but using a triangle shape arena of similar size to the home cage with one of the corners having a 45-degree angle where the cotton swab was placed. A transparent plexiglass wall would allow viewing of the pupil when mice explored the cotton swab at close proximity. Images of the pupil where captured during the first 10-second bouts of exploration.

#### Two-Photon Calcium Imaging in Head-fixed Mice

A two-photon microscope (FVMPE-RS, Olympus, Center Valley, PA) was used to visualize activity dynamics of *Pnoc*^+^ neurons in BNST *in vivo* in head-fixed mice while they underwent odor exposure with pupillometry. A virus encoding the Cre-dependent calcium indicator GCaMP6s (AAVdj-EF1α-DIO-GCaMP6s; 3.1 x 10^12^ infectious units/mL) was injected into BNST of *Pnoc-Cre* mice (see Surgery and histology section). After a minimum of 8 weeks to allow sufficient time for virus transport and infection, mice underwent the head-fixed freely moving odor exposure assay described above, during which GCaMP6s-expressing neurons were visualized using two-photon microscopy.

The two-photon microscope used was equipped with the following to allow imaging of BNST *in vivo*: a hybrid scanning core set with galvanometers and fast scan resonant scanners (which allows up to 30 Hz frame -rate acquisition; set at 5 Hz), GaAsP-PMT photo detectors with adjustable voltage, gain, and offset features, a single green/red NDD filter cube, a long working distance 20x objective (air) designed for optical transmission at infrared wavelengths (LCPLN20XIR, 0.45 NA, 8.3 mm WD, Olympus, Center Valley, PA), a software-controlled modular *xy* stage loaded on a manual *z*-deck, and a tunable Mai-Tai Deep See laser system (laser set to 955 nm, ∼100 fs pulse width, Spectra Physics, Santa Clara, CA) with automated four-axis alignment. Prior to testing, the optimal field of view (FOV) was selected by adjusting the imaging plane (*z*-axis). Two-photon scanning was triggered by an Arduino microcontroller and video was collected for each testing epoch (baseline, water or odor). Data were both acquired and processed using FluoView FV1200 and CellSens software packages (Olympus, Center Valley, PA). Following data acquisition, videos were motion corrected using a planar hidden Markov model (SIMA v1.3) (Kaifosh et al., 2014), calcium transients and deconvolved events were extracted from individual ROI’s using constraint non-negative matrix factorization algorithms (CNMF) (Zhou et al., 2018) and data was analyzed using custom data analysis pipelines written in Python (see Quantification and Statistical Analysis section).

#### Head-fixed Odor Swab Exposure with Pupillometry

For exposing odors in head-fixed freely moving mice, experimental events and behavioral recordings were orchestrated using custom-designed hardware interfaced with microcontrollers (Arduino) and Python using custom code. Odor delivery relied on a custom-made conveyor system that carried a cotton swab with odor source along a 25-cm track over 6 s to and from the animal. The cotton swab remained in close proximity to the animal for a 10-second bout. We assessed locomotor activity of head-fixed animals using a custom-made running disc. The disc was fixed under the head-fixed animal, which allowed movement similar to a rodent flying saucer wheel. Rotational changes were measured by a rotary encoder (Sparkfun, Boulder, CO) every 50 ms without regard to direction of rotation. Pupil recordings were made using a monochromatic CMOS camera with macro zoom lens (MVL7000 & DCC1545M, ThorLabs, Newton, NJ) at 10 frames per second. An infrared light (Thorlabs, Newton, NJ) was used to illuminate the eye in optogenetic experiments. For two-photon experiments, the illumination light from the objective was sufficient to visualize the eye (here the light transmitted through nervous tissue and out the pupil, thus the pupil was brighter than the cornea). An ultraviolet light (Thorlabs, Newton, NJ) was used to adjust the pupil size to avoid a ceiling or floor effect of pupil changes as necessary.

Experimentation began after minimal pupillary responses were observed to the approaching of a dry cotton swab (6 days). The day of testing, mice were exposed to 3 epochs (5 minutes each) that consisted of 5 baseline bouts (dry cotton swab), 5 control bouts (cotton swab with 2.5 μl of distilled H_2_O), and 5 odor bouts (cotton swab with wither 2.5 μl of TMT or 2.5 μl of Peanut oil). The first 2 bouts of each epoch were used for analysis to assess responses. A low dose of TMT was used to minimize freezing responses and maintain ambulation.

Pupil changes were assessed offline after experimentation. A median filter was applied to each pupil recording frame before pupil diameter was measured. We used OpenCV to identify the pupil within each frame and morphological processing (erosion and dilation) to further filter noise from the image. The diameter of the pupil was measured by fitting a bounding box, and the length of its horizontal sides were used as the pupil diameter since this also measured pupil diameter fairly well during mid-blink. Calculated diameter measures were then filtered using a rolling 1-s median filter.

#### Optogenetics

Optogenetic experiments were performed as previously described (Sparta et al., 2011). Briefly, a virus encoding the Cre-inducible channel-rhodopsin-2 (AAV5-ef1α-DIO-hChR2(H134R)-eYFP; 5.0 x 10^12^ infectious units per ml) was injected into BNST of either *Pnoc-Cre* mice or their wild type littermates as controls. For photoactivation manipulations in ChR2 or control mice, the laser (473 nm; 8–10 mW) was turned on for 5-ms pulses (20 Hz) during a 3 min period, followed or preceded by 3 min periods were the laser was off. All mice were habituated to the tether for 3 days prior to behavioral testing. Following behavioral experiments, histological verification of fluorescence and optical fiber placement were performed.

#### Real-Time Place Preference

Mice were placed into a rectangular two-compartment arena (52.5 x 25.5 x 25.5 cm) as previously described (Jennings et al., 2013a). Mice were allowed to freely explore the arena for 20 min. Entry into one of the compartments triggered constant 20 Hz photostimulation (473 nm; 8–10 mW). Entry into the other chamber ended the photostimulation. The side paired with photostimulation was counterbalanced across mice. Time spent in the stimulation side was calculated using automated tracking software (Ethovision XT 11, Noldus, Leesburg, VA).

#### Head-fixed Stationary Assay with Pupillometry

Mice were head-fixed as previously described (Otis et al., 2017). Physiological and licking measures were obtained using a custom designed apparatus. A piezo sensor under the mouse monitored general movement in the tube. A pulse oximeter placed near the neck was used to measure heart rate. Mice received unpredictable drops of sucrose (10% in water, 2.0-2.5 μl, ∼1 drop/min) for 30 min using a gravity-driven solenoid through a ∼18-G steel tube. Mice were habituated to the setup for 6 days. Measurements were recorded using a LabJack data acquisition box (U12 Series, LabJack Corp., Lakewood, CO). Once mice habituated to the apparatus, as evident by a reduced heart rate as compared to Day 1 (6 days), optogenetic experiments were performed while pupil videos, movement (piezo sensor) and heart rate (pulse oximeter) was tracked with an Arduino microcontroller and recorded with custom software (written in Python) during a single laser off (3 min) and laser on (3 min) period.

#### Tissue Isolation and Single-cell cDNA Library Preparation

Mice were anesthetized with 390 g/kg sodium pentobarbital, 500 mg/kg phenytoin sodium and transcardially perfused with 20 mL in ice-cold sodium-substituted aCSF (NMDG-aCSF: 96 mM NMDG, 2.5 mM KCl, 1.35 mM NaH_2_PO_4_, 30 mM NaHCO_3_, 20 mM HEPES, 25 mM glucose, 2 mM thiourea, 5 mM Na^+^ascorbate, 3 mM Na^+^pyruvate, 0.6 mM glutathione-ethyl-ester, 2 mM N-acetyl-cysteine, 0.5 mM CaCl_2_, 10 mM MgSO_4_; pH 7.35–7.40, 300-305 mOsm) modified from (Ting et al., 2014). Brains were isolated and three 300 μm sections beginning at ∼0.45 mm Bregma were collected in ice-cold NMDG-aCSF on a vibratome (Leica, VT 1200, Germany). Sections from 6 mice at a time (total of 4 batches with 24 mice) were recovered in NMDG-aCSF supplemented with 500 nM TTX, 10 μM APV, 10 μM DNQX (NMDG-aCSF-R) for 15 minutes after the addition of the last slice. The BNST was then isolated with 0.75 and 0.50 mm Palkovitz punches and digested in NMDG-aCSF-R containing 1.0 mg/mL pronase for 30 minutes at room temperature. Tissue was then triturated with a patch pipet fire-polished to an internal diameter of 300 μm in 1.0 mL of NMDG-aCSF-R supplemented with 0.05% BSA (NMDG-aCSF-BSA) to dissociate. The suspension transferred to 12 mL NMDG-aCSF-BSA and sedimented at 220 x *g* for 6 minutes at 18°C to wash. The supernatant was removed, and cells were resuspended in 1 mL NMDG-aCSF-BSA. To fix the cells (Alles et al., 2017), 4.0 mL of ice-cold 100% methanol was added dropwise to the suspension while gently swirling the tube. Cells were then incubated for 30 minutes on ice and transferred to −80 °C. To rehydrate suspensions prior to Drop-seq, cells were removed from −80 °C and incubated on ice for 15 minutes. Cells were then sedimented at 500 x *g* for 5 minutes at 4 °C, resuspended in 5 mL of PBS supplemented with 0.01% BSA (PBS-BSA), and incubated for 5 minutes on ice. The suspension was then sedimented at 220 x *g* for 6 minutes at 18°C and resuspended in 1.0 mL of PBS-BSA for a final concentration of ∼2.6-3.2 x 10^5^ cells/mL. Rehydration and droplet generation was performed on fixed samples within 3 weeks of fixation.

Drop-seq was performed as previously described in with modifications (Macosko et al., 2015). Single-cell capture was performed on a glass microfluidics device (Dolomite Microfluidics, United Kingdom) with aqueous flow at 40 μL/min and oil at 200 μL/min. Beads were loaded at ∼200 beads/μL. Reverse transcription, ExoI digestion, and PCR were performed as previously described, but with 11 cycles for second stage of amplification. PCR products were pooled by batch, purified on SPRI beads (Axygen, Union City, CA), and indexed using Nextera XT with 800 *p*g input per batch. Purified tagmentation products were pooled by mass according to the estimated number of cells per pool member as quantified by a Qubit dsDNA HS Assay. Sequencing was performed at the UNC High Throughput Sequencing Facility on a lllumina HiSeq2500 using Paired-End 2×50 Rapid Run v2 chemistry.

### QUANTIFICATION AND STATISTICAL ANALYSIS

#### Behavioral Optogenetics and Electrophysiology Data Analysis

For data obtained from the optogenetic and patch-clamp electrophysiology experiments, data were analyzed using Prism 7 (GraphPad Sotware Inc., La Jolla, CA). Mean values are accompanied by SEM values. Comparisons were tested using paired or unpaired t-tests. Two-way ANOVA tests followed by either Tukey’s post-hoc tests or Bonferroni post-hoc comparisons were applied for comparisons with more than two groups, n.s. p > 0.05, *p < 0.05, **p < 0.01, ***p < 0.001.

#### Calcium Imaging Analysis

Calcium imaging recordings were first motion corrected using a planar hidden Markov model (Kaifosh et al., 2014). Neurons were identified, and their calcium signals were extracted using a modified version of constrained nonnegative matrix factorization (CNMF) (Zhou et al., 2018), allowing us to segregate spatially overlapping signals. This extracted signal was adjusted (scaled) to account for variations in fluorescence intensities among cells by the standard deviation of a neuron’s fluorescence throughout the Control odor exposure. For head-fixed, odor-presentation experiments, neuronal activity was aligned to the presentation of the odor. Neurons were classified as excitatory or inhibitory to proximity of TMT or Peanut oil if the fluorescence values for frames differed between near and far location—defined using a Mann-Whitney U test with Bonferroni correction. Correlations in activity and behavior were calculated using the Spearman correlation coefficient.

#### Single-Cell Sequencing Clustering and Analysis

Demultiplexing was performed allowing 1 mismatch with Illumina bcl2fastq v2.18.0.12. Initial processing and generation of digital expression matrices was performed with Drop-seq_tools v1.12 and Picard Tools v2.2.4 (Macosko et al., 2015). Alignment was performed using STAR v2.4.2a with 72 GB of RAM and 16 threads. Clustering was performed in R using Seurat v1.4.0.16 unless otherwise noted. Prior to clustering, cells were filtered by ≥ 500 unique genes, ≤ 20,000 unique molecules, and ≤ 5 percent mitochondrial reads. Filtered data was scaled to the median number of unique molecules and log(x+1) transformed. Zero-variance genes were removed from the data, and batch correction was performed with ComBat (Johnson et al., 2007) from SVA v3.220 (Leek et al., 2012) using parametric adjustment on a model matrix containing number of unique genes and molecules, and percent mitochondrial reads. Four batches were included, each containing six animals that were pooled during tissue isolation. Relative log expression by cell and mean expression correlation across batches were used to assess the correction. Only genes detected in all batches were included in the analysis.

Variable genes were selected with a cutoff of 0.5 standard deviations from the mean dispersion within a bin (Macosko et al., 2015). Variable genes were used as the basis for principal components analysis, and cluster calling was performed on principal components using the Louvain algorithm with multilevel refinement under default settings. Principal components were reduced and visualized via t-distributed stochastic neighbor embedding (tSNE) using the first 20 components and a resolution of 1.3. Clusters were reordered on a hierarchically-clustered distance matrix based on all genes. Features were identified using a single-cell likelihood-ratio test^6^ implemented in Seurat. To identify cluster-specific features, genes in each cluster were tested against those in either the nearest cluster or node in the hierarchically-clustered dendrogram. Analysis from pre-processing to digital expression matrices were run on a Dell blade-based cluster running RedHat Enterprise Linux 5.6. Cluster calling and tSNE were run on a similar cluster running RedHat Enterprise Linux 7.3. All other steps were run on macOS 10.13.3.

### DATA AND SOFTWARE AVAILABILITY

Code used for analysis are openly available online (https://github.com/stuberlab). Single cell sequencing data is available at GEO (accession GSE132067). All other data are available upon request from the corresponding author.

## REFERENCES

Ahrens, S., Wu, M.V., Furlan, A., Hwang, G.-R., Paik, R., Li, H., Penzo, M.A., Tollkuhn, J., and Li, B. (2018). A Central Extended Amygdala Circuit That Modulates Anxiety. J. Neurosci. 38, 5567–5583.

Alles, J., Karaiskos, N., Praktiknjo, S.D., Grosswendt, S., Wahle, P., Ruffault, P.-L., Ayoub, S., Schreyer, L., Boltengagen, A., Birchmeier, C., et al. (2017). Cell fixation and preservation for droplet-based single-cell transcriptomics. BMC Biol. 15, 44.

Boom, A., Mollereau, C., Meunier, J.C., Vassart, G., Parmentier, M., Vanderhaeghen, J.J., and Schiffmann, S.N. (1999). Distribution of the nociceptin and nocistatin precursor transcript in the mouse central nervous system. Neuroscience 91, 991–1007.

Calhoon, G.G., and Tye, K.M. (2015). Resolving the neural circuits of anxiety. Nat. Neurosci. 18, 1394–1404.

Cascardi, M., Armstrong, D., Chung, L., and Paré, D. (2015). Pupil Response to Threat in Trauma-Exposed Individuals With or Without PTSD. J Trauma Stress 28, 370–374.

Craske, M.G., Rauch, S.L., Ursano, R., Prenoveau, J., Pine, D.S., and Zinbarg, R.E. (2009). What is an anxiety disorder? Depress Anxiety 26, 1066–1085.

Crowley, N.A., Bloodgood, D.W., Hardaway, J.A., Kendra, A.M., McCall, J.G., Al-Hasani, R., McCall, N.M., Yu, W., Schools, Z.L., Krashes, M.J., et al. (2016). Dynorphin Controls the Gain of an Amygdalar Anxiety Circuit. Cell Rep 14, 2774–2783.

Dabrowska, J., Hazra, R., Ahern, T.H., Guo, J.-D., McDonald, A.J., Mascagni, F., Muller, J.F., Young, L.J., and Rainnie, D.G. (2011). Neuroanatomical evidence for reciprocal regulation of the corticotrophin-releasing factor and oxytocin systems in the hypothalamus and the bed nucleus of the stria terminalis of the rat: Implications for balancing stress and affect. Psychoneuroendocrinology 36, 1312–1326.

Dong, H.W., Petrovich, G.D., and Swanson, L.W. (2001). Topography of projections from amygdala to bed nuclei of the stria terminalis. Brain Res. Brain Res. Rev. 38, 192–246.

Duvarci, S., Bauer, E.P., and Paré, D. (2009). The bed nucleus of the stria terminalis mediates inter-individual variations in anxiety and fear. J. Neurosci. 29, 10357–10361.

Gafford, G.M., and Ressler, K.J. (2015). GABA and NMDA receptors in CRF neurons have opposing effects in fear acquisition and anxiety in central amygdala vs. bed nucleus of the stria terminalis. Horm Behav 76, 136–142.

Ghosh, K.K., Burns, L.D., Cocker, E.D., Nimmerjahn, A., Ziv, Y., Gamal, A.E., and Schnitzer, M.J. (2011). Miniaturized integration of a fluorescence microscope. Nat. Methods 8, 871–878.

Giardino, W.J., Eban-Rothschild, A., Christoffel, D.J., Li, S.-B., Malenka, R.C., and de Lecea, L. (2018). Parallel circuits from the bed nuclei of stria terminalis to the lateral hypothalamus drive opposing emotional states. Nat. Neurosci.

Goodson, J.L., and Wang, Y. (2006). Valence-sensitive neurons exhibit divergent functional profiles in gregarious and asocial species. Proc. Natl. Acad. Sci. U.S.A. 103, 17013–17017.

Gungor, N.Z., and Paré, D. (2016). Functional Heterogeneity in the Bed Nucleus of the Stria Terminalis. J. Neurosci. 36, 8038–8049.

Hammack, S.E., Guo, J.-D., Hazra, R., Dabrowska, J., Myers, K.M., and Rainnie, D.G. (2009). The response of neurons in the bed nucleus of the stria terminalis to serotonin: implications for anxiety. Prog. Neuropsychopharmacol. Biol. Psychiatry 33, 1309–1320.

Hardaway, J.A., Halladay, L.R., Mazzone, C.M., Pati, D., Bloodgood, D.W., Kim, M., Jensen, J., DiBerto, J.F., Boyt, K.M., Shiddapur, A., et al. (2019). Central Amygdala Prepronociceptin-Expressing Neurons Mediate Palatable Food Consumption and Reward. Neuron.

Harris, A.Z., and Gordon, J.A. (2015). Long-range neural synchrony in behavior. Annu. Rev. Neurosci. 38, 171–194.

Hermans, E.J., Henckens, M.J.A.G., Roelofs, K., and Fernández, G. (2013). Fear bradycardia and activation of the human periaqueductal grey. Neuroimage 66, 278–287.

Ikeda, K., Watanabe, M., Ichikawa, T., Kobayashi, T., Yano, R., and Kumanishi, T. (1998). Distribution of prepro-nociceptin/orphanin FQ mRNA and its receptor mRNA in developing and adult mouse central nervous systems. J. Comp. Neurol. 399, 139–151.

Jennings, J.H., Sparta, D.R., Stamatakis, A.M., Ung, R.L., Pleil, K.E., Kash, T.L., and Stuber, G.D. (2013a). Distinct extended amygdala circuits for divergent motivational states. Nature 496, 224–228.

Jennings, J.H., Rizzi, G., Stamatakis, A.M., Ung, R.L., and Stuber, G.D. (2013b). The Inhibitory Circuit Architecture of the Lateral Hypothalamus Orchestrates Feeding. Science 341, 1517– 1521.

Johnson, W.E., Li, C., and Rabinovic, A. (2007). Adjusting batch effects in microarray expression data using empirical Bayes methods. Biostatistics 8, 118–127.

Kaifosh, P., Zaremba, J.D., Danielson, N.B., and Losonczy, A. (2014). SIMA: Python software for analysis of dynamic fluorescence imaging data. Front Neuroinform 8, 80.

Kash, T.L., Pleil, K.E., Marcinkiewcz, C.A., Lowery-Gionta, E.G., Crowley, N., Mazzone, C., Sugam, J., Hardaway, J.A., and McElligott, Z.A. (2015). Neuropeptide regulation of signaling and behavior in the BNST. Mol. Cells 38, 1–13.

Kim, S.-Y., Adhikari, A., Lee, S.Y., Marshel, J.H., Kim, C.K., Mallory, C.S., Lo, M., Pak, S., Mattis, J., Lim, B.K., et al. (2013). Diverging neural pathways assemble a behavioural state from separable features in anxiety. Nature 496, 219–223.

Kodani, S., Soya, S., and Sakurai, T. (2017). Excitation of GABAergic Neurons in the Bed Nucleus of the Stria Terminalis Triggers Immediate Transition from Non-Rapid Eye Movement Sleep to Wakefulness in Mice. J. Neurosci. 37, 7164–7176.

Koob, G.F., and Heinrichs, S.C. (1999). A role for corticotropin releasing factor and urocortin in behavioral responses to stressors. Brain Res. 848, 141–152.

Lang, P.J., and McTeague, L.M. (2009). The anxiety disorder spectrum: fear imagery, physiological reactivity, and differential diagnosis. Anxiety Stress Coping 22, 5–25.

Lebow, M.A., and Chen, A. (2016). Overshadowed by the amygdala: the bed nucleus of the stria terminalis emerges as key to psychiatric disorders. Mol. Psychiatry 21, 450–463.

de Lecea, L., Carter, M.E., and Adamantidis, A. (2012). Shining light on wakefulness and arousal. Biol. Psychiatry 71, 1046–1052.

LeDoux, J.E., and Pine, D.S. (2016). Using Neuroscience to Help Understand Fear and Anxiety: A Two-System Framework. Am J Psychiatry 173, 1083–1093.

Leek, J.T., Johnson, W.E., Parker, H.S., Jaffe, A.E., and Storey, J.D. (2012). The sva package for removing batch effects and other unwanted variation in high-throughput experiments. Bioinformatics 28, 882–883.

Lei, K., Cushing, B.S., Musatov, S., Ogawa, S., and Kramer, K.M. (2010). Estrogen receptor-alpha in the bed nucleus of the stria terminalis regulates social affiliation in male prairie voles (Microtus ochrogaster). PLoS ONE 5, e8931.

Li, Y., Mathis, A., Grewe, B.F., Osterhout, J.A., Ahanonu, B., Schnitzer, M.J., Murthy, V.N., and Dulac, C. (2017). Neuronal Representation of Social Information in the Medial Amygdala of Awake Behaving Mice. Cell 171, 1176–1190.e17.

Macosko, E.Z., Basu, A., Satija, R., Nemesh, J., Shekhar, K., Goldman, M., Tirosh, I., Bialas, A.R., Kamitaki, N., Martersteck, E.M., et al. (2015). Highly Parallel Genome-wide Expression Profiling of Individual Cells Using Nanoliter Droplets. Cell 161, 1202–1214.

Marcinkiewcz, C.A., Mazzone, C.M., D’Agostino, G., Halladay, L.R., Hardaway, J.A., DiBerto, J.F., Navarro, M., Burnham, N., Cristiano, C., Dorrier, C.E., et al. (2016). Serotonin engages an anxiety and fear-promoting circuit in the extended amygdala. Nature 537, 97–101.

Marton, T.F., and Sohal, V.S. (2016). Of Mice, Men, and Microbial Opsins: How Optogenetics Can Help Hone Mouse Models of Mental Illness. Biol. Psychiatry 79, 47–52.

Mazzone, C.M., Pati, D., Michaelides, M., DiBerto, J., Fox, J.H., Tipton, G., Anderson, C., Duffy, K., McKlveen, J.M., Hardaway, J.A., et al. (2018). Acute engagement of G_q_-mediated signaling in the bed nucleus of the stria terminalis induces anxiety-like behavior. Molecular Psychiatry 23, 143–153.

McDonald, A.J. (1989). Coexistence of somatostatin with neuropeptide Y, but not with cholecystokinin or vasoactive intestinal peptide, in neurons of the rat amygdala. Brain Res. 500, 37–45.

McElligott, Z.A., and Winder, D.G. (2009). Modulation of glutamatergic synaptic transmission in the bed nucleus of the stria terminalis. Prog. Neuropsychopharmacol. Biol. Psychiatry 33, 1329– 1335.

McHenry, J.A., Otis, J.M., Rossi, M.A., Robinson, J.E., Kosyk, O., Miller, N.W., McElligott, Z.A., Budygin, E.A., Rubinow, D.R., and Stuber, G.D. (2017). Hormonal gain control of a medial preoptic area social reward circuit. Nat Neurosci 20, 449–458.

Namboodiri, V.M.K., Otis, J.M., van Heeswijk, K., Voets, E.S., Alghorazi, R.A., Rodriguez-Romaguera, J., Mihalas, S., and Stuber, G.D. (2019). Single-cell activity tracking reveals that orbitofrontal neurons acquire and maintain a long-term memory to guide behavioral adaptation. Nature Neuroscience.

Neal, C.R., Mansour, A., Reinscheid, R., Nothacker, H.P., Civelli, O., and Watson, S.J. (1999). Localization of orphanin FQ (nociceptin) peptide and messenger RNA in the central nervous system of the rat. J. Comp. Neurol. 406, 503–547.

Newman Sarah Winans (2006). The Medial Extended Amygdala in Male Reproductive Behavior A Node in the Mammalian Social Behavior Network. Annals of the New York Academy of Sciences 877, 242–257.

Nguyen, A.Q., Cruz, J.A.D.D., Sun, Y., Holmes, T.C., and Xu, X. (2016). Genetic cell targeting uncovers specific neuronal types and distinct subregions in the bed nucleus of the stria terminalis. J Comp Neurol 524, 2379–2399.

Otis, J.M., Namboodiri, V.M.K., Matan, A.M., Voets, E.S., Mohorn, E.P., Kosyk, O., McHenry, J.A., Robinson, J.E., Resendez, S.L., Rossi, M.A., et al. (2017). Prefrontal cortex output circuits guide reward seeking through divergent cue encoding. Nature 543, 103–107.

Parker, K.E., Pedersen, C.E., Gomez, A.M., Spangler, S.M., Walicki, M.C., Feng, S.Y., Stewart, S.L., Otis, J.M., Al-Hasani, R., McCall, J.G., et al. (2019). A Paranigral VTA Nociceptin Circuit that Constrains Motivation for Reward. Cell 178, 653–671.e19.

Patriquin, M.A., Hartwig, E.M., Friedman, B.H., Porges, S.W., and Scarpa, A. (2019). Autonomic response in autism spectrum disorder: Relationship to social and cognitive functioning. Biol Psychol 145, 185–197.

Perusini, J.N., and Fanselow, M.S. (2015). Neurobehavioral perspectives on the distinction between fear and anxiety. Learn. Mem. 22, 417–425.

Pompolo, S., Ischenko, O., Pereira, A., Iqbal, J., and Clarke, I.J. (2005). Evidence that projections from the bed nucleus of the stria terminalis and from the lateral and medial regions of the preoptic area provide input to gonadotropin releasing hormone (GNRH) neurons in the female sheep brain. Neuroscience 132, 421–436.

Price, R.B., Siegle, G.J., Silk, J.S., Ladouceur, C., McFarland, A., Dahl, R.E., and Ryan, N.D. (2013). Sustained neural alterations in anxious youth performing an attentional bias task: a pupilometry study. Depress Anxiety 30, 22–30.

Reimer, J., Froudarakis, E., Cadwell, C.R., Yatsenko, D., Denfield, G.H., and Tolias, A.S. (2014). Pupil fluctuations track fast switching of cortical states during quiet wakefulness. Neuron 84, 355–362.

Resendez, S.L., Jennings, J.H., Ung, R.L., Namboodiri, V.M.K., Zhou, Z.C., Otis, J.M., Nomura, H., McHenry, J.A., Kosyk, O., and Stuber, G.D. (2016). Visualization of cortical, subcortical and deep brain neural circuit dynamics during naturalistic mammalian behavior with head-mounted microscopes and chronically implanted lenses. Nat Protoc 11, 566–597.

Root, C.M., Denny, C.A., Hen, R., and Axel, R. (2014). The participation of cortical amygdala in innate, odour-driven behaviour. Nature 515, 269–273.

Schmidt, F.M., Sander, C., Dietz, M.-E., Nowak, C., Schröder, T., Mergl, R., Schönknecht, P., Himmerich, H., and Hegerl, U. (2017). Brain arousal regulation as response predictor for antidepressant therapy in major depression. Sci Rep 7, 45187.

Schneider, M., Leuchs, L., Czisch, M., Sämann, P.G., and Spoormaker, V.I. (2018). Disentangling reward anticipation with simultaneous pupillometry / fMRI. Neuroimage 178, 11– 22.

Shin, L.M., and Liberzon, I. (2010). The neurocircuitry of fear, stress, and anxiety disorders. Neuropsychopharmacology 35, 169–191.

Singewald, N., Salchner, P., and Sharp, T. (2003). Induction of c-Fos expression in specific areas of the fear circuitry in rat forebrain by anxiogenic drugs. Biol. Psychiatry 53, 275–283.

Sparta, D.R., Stamatakis, A.M., Phillips, J.L., Hovelsø, N., van Zessen, R., and Stuber, G.D. (2011). Construction of implantable optical fibers for long-term optogenetic manipulation of neural circuits. Nat Protoc 7, 12–23.

Sparta, D.R., Jennings, J.H., Ung, R.L., and Stuber, G.D. (2013). Optogenetic strategies to investigate neural circuitry engaged by stress. Behav. Brain Res. 255, 19–25.

Stamatakis, A.M., Sparta, D.R., Jennings, J.H., McElligott, Z.A., Decot, H., and Stuber, G.D. (2014). Amygdala and bed nucleus of the stria terminalis circuitry: Implications for addiction-related behaviors. Neuropharmacology 76 Pt B, 320–328.

Straube, T., Mentzel, H.-J., and Miltner, W.H.R. (2007). Waiting for spiders: brain activation during anticipatory anxiety in spider phobics. Neuroimage 37, 1427–1436.

Stuber, G.D., and Wise, R.A. (2016). Lateral hypothalamic circuits for feeding and reward. Nat. Neurosci. 19, 198–205.

Ting, J.T., Daigle, T.L., Chen, Q., and Feng, G. (2014). Acute brain slice methods for adult and aging animals: application of targeted patch clamp analysis and optogenetics. Methods Mol. Biol. 1183, 221–242.

Touriño, C., Eban-Rothschild, A., and de Lecea, L. (2013). Optogenetics in psychiatric diseases. Curr. Opin. Neurobiol. 23, 430–435.

Tovote, P., Fadok, J.P., and Lüthi, A. (2015). Neuronal circuits for fear and anxiety. Nat. Rev. Neurosci. 16, 317–331.

Tummeltshammer, K., Feldman, E.C.H., and Amso, D. (2019). Using pupil dilation, eye-blink rate, and the value of mother to investigate reward learning mechanisms in infancy. Dev Cogn Neurosci 36, 100608.

Urbano, F.J., Bisagno, V., and Garcia-Rill, E. (2017). Arousal and drug abuse. Behav. Brain Res. 333, 276–281.

Walker, D.L., Miles, L.A., and Davis, M. (2009). Selective participation of the bed nucleus of the stria terminalis and CRF in sustained anxiety-like versus phasic fear-like responses. Prog. Neuropsychopharmacol. Biol. Psychiatry 33, 1291–1308.

Wilhelm, F.H., and Roth, W.T. (2001). The somatic symptom paradox in DSM-IV anxiety disorders: suggestions for a clinical focus in psychophysiology. Biol Psychol 57, 105–140.

Yassa, M.A., Hazlett, R.L., Stark, C.E.L., and Hoehn-Saric, R. (2012). Functional MRI of the amygdala and bed nucleus of the stria terminalis during conditions of uncertainty in generalized anxiety disorder. J Psychiatr Res 46, 1045–1052.

Zhou, P., Resendez, S.L., Rodriguez-Romaguera, J., Jimenez, J.C., Neufeld, S.Q., Giovannucci, A., Friedrich, J., Pnevmatikakis, E.A., Stuber, G.D., Hen, R., et al. (2018). Efficient and accurate extraction of in vivo calcium signals from microendoscopic video data. ELife Sciences 7, e28728.

